# Tumor Heterogeneity Induces Pro- and Anti-metastatic Myeloid Cell Phenotypes in Breast Cancer Lung Metastasis

**DOI:** 10.64898/2026.03.27.714678

**Authors:** Daphne A Superville, Eva Chrenková, Yimin Zheng, Alexis J. Combes, Iros Barozzi, Zena Werb, André F. Rendeiro, Christopher S. McGinnis, Andrei Goga, Juliane Winkler

## Abstract

Metastasis is a major cause of cancer-related mortality, yet targeting metastatic cells directly has been largely unsuccessful due to their plasticity and heterogeneity. Myeloid cells play diverse pro- and anti-metastatic functions and are an attractive alternative target for treating metastasis, but how tumor heterogeneity influences myeloid cell phenotypes during metastasis remains poorly understood. Here, we profiled myeloid cells from primary tumors and matched metastatic lungs of 12 heterogeneous and differentially metastatic patient-derived xenograft models of breast cancer. Comparative analysis of cell type abundances revealed distinct myeloid remodeling specific to primary tumors or metastatic lungs. Beyond compositional differences, we identified gene expression programs that were associated with metastatic burden, such as number or size of metastatic nodules, indicating distinct microenvironmental requirements for metastatic seeding and outgrowth. Examining these metastasis-associated programs using time-course datasets, we discovered an evolution from anti- to pro-metastatic monocyte phenotypes during metastatic progression. We further showed that this phenotypic shift was driven by an increase in two distinct myeloid-derived suppressor cell signatures, and a transcriptionally regulated impairment of monocyte differentiation leading to the depletion of non-classical monocytes. Our results comprehensively dissect the heterogeneity of myeloid cell phenotypes across primary tumor and metastatic sites, opening novel avenues for myeloid-targeting therapies specific to metastasis.

## Introduction

Breast cancer (BC) is the leading cause of cancer deaths in women worldwide [1]. While 5-year survival rates have improved for localized and regional disease (87– 99%), metastatic BC survival rates remain relatively low (33%) [2]. Unlike primary tumorigenesis, since metastasis is largely not driven by any specific genetic alterations, developing targeted treatments focusing on tumor cells has proven difficult [3, 4]. Therefore, there is an urgent clinical need to develop alternative therapeutic strategies to treat or prevent metastasis.

Innate immune cells are key regulators of metastasis, playing both pro- and anti-metastatic roles. For example, myeloid-derived suppressor cells (MDSCs) are a subset of suppressive myeloid cells of granulocytic or monocytic origin [5]. A range of pro-metastatic functions have been attributed to MDSCs, including promoting tumor cell invasiveness, suppressing adaptive immune cells, and promoting tumor cell outgrowth at the metastatic site [6,7]. Conversely, (non-MDSC) subsets of neutrophils and monocytes can suppress metastasis. For example, IFN-responsive monocytes have been shown to recruit neutrophils that kill tumor cells in the metastatic lung of BC models [8]. Additionally, non-classical monocytes can recruit NK cells to eliminate transiting tumor cells in the vasculature [9, 10]. These diverse myeloid cell phenotypes and their complex remodelling during metastatic progression remain incompletely understood, challenging the development of myeloid cell-targeted immunotherapies for metastasis. We and others have profiled the immune metastatic lung niche of BC, revealing dynamic changes in myeloid cell composition and phenotypes during metastatic progression [11–14]. However, current studies lack matching samples for primary tumors and metastases for comparative analysis, or use mouse models, missing the effect of human tumor heterogeneity on myeloid remodeling [11, 13]. Further, limited human tissue availability and the prolonged, variable time to the clinical onset of metastasis make analysis of matched samples nearly impossible. Therefore, heterogeneous and human-relevant tumor models are needed to disentangle the diversity of innate immune cell behaviors underlying different metastatic phenotypes.

To explore how BC heterogeneity shapes myeloid remodeling during metastatic progression, we performed single-cell RNA sequencing on immune cells isolated from matched primary tumors and metastases of BC patient-derived xenograft (PDX) models with varying propensities to metastasize to the lung [15]. Our findings reveal distinct myeloid microenvironments in primary tumors and metastatic lungs, and transcriptional diversity associated with metastatic progression. Overall, this study elucidates the evolution of myeloid phenotypes within the metastatic niche and highlights potential targets for metastasis-specific myeloid therapies.

## Results

### Single-cell profiling of myeloid cells in primary and metastatic breast cancers

To dissect the heterogeneity and evolution of myeloid cell phenotypes during metastatic progression in BC, we investigated the immune landscape of matched primary tumors and their corresponding lung metastases of 12 PDX models. We previously showed that myeloid cells are substantially remodeled during metastatic progression in one highly metastatic PDX model [12]. Despite the lack of T, B, and NK cells and species discrepancy, PDXs serve as suitable models for investigating the impact of tumor heterogeneity on myeloid remodeling in the metastatic niche within a controlled genetic background. Lungs and mammary glands from non-tumor bearing mice served as controls. PDX models included luminal B and triple-negative subtypes characterized by high intra- and inter-tumoral transcriptional heterogeneity and varying metastatic potentials [15]. PDX models were previously categorized as models with either high (4 models), moderate (3 models), or low (5 models) metastatic potential based on histological characterization evaluating the number and size of metastatic nodules and metastatic burden (percent metastatic area) in lung tissue [15]. Primary tumors, metastatic lungs, and control tissues were collected, digested into a single cell suspension, MULTI-seq barcoded [12] and sorted for CD45^+^ immune cells prior to pooling and sequencing using the 10X Genomics platform (Fig. 1A). After sample demultiplexing and species-aware QC, our dataset consisted of 56,931 cells sequenced across six batches, representing 52 samples of 12 PDX models and 16 samples of 2 control tissues. Each PDX was represented by at least two biological replicates for each tissue, with tumor/mammary gland samples averaging 543 cells and lung samples averaging 1,114 cells (Supp. Fig. S1A).

**Figure 1.**
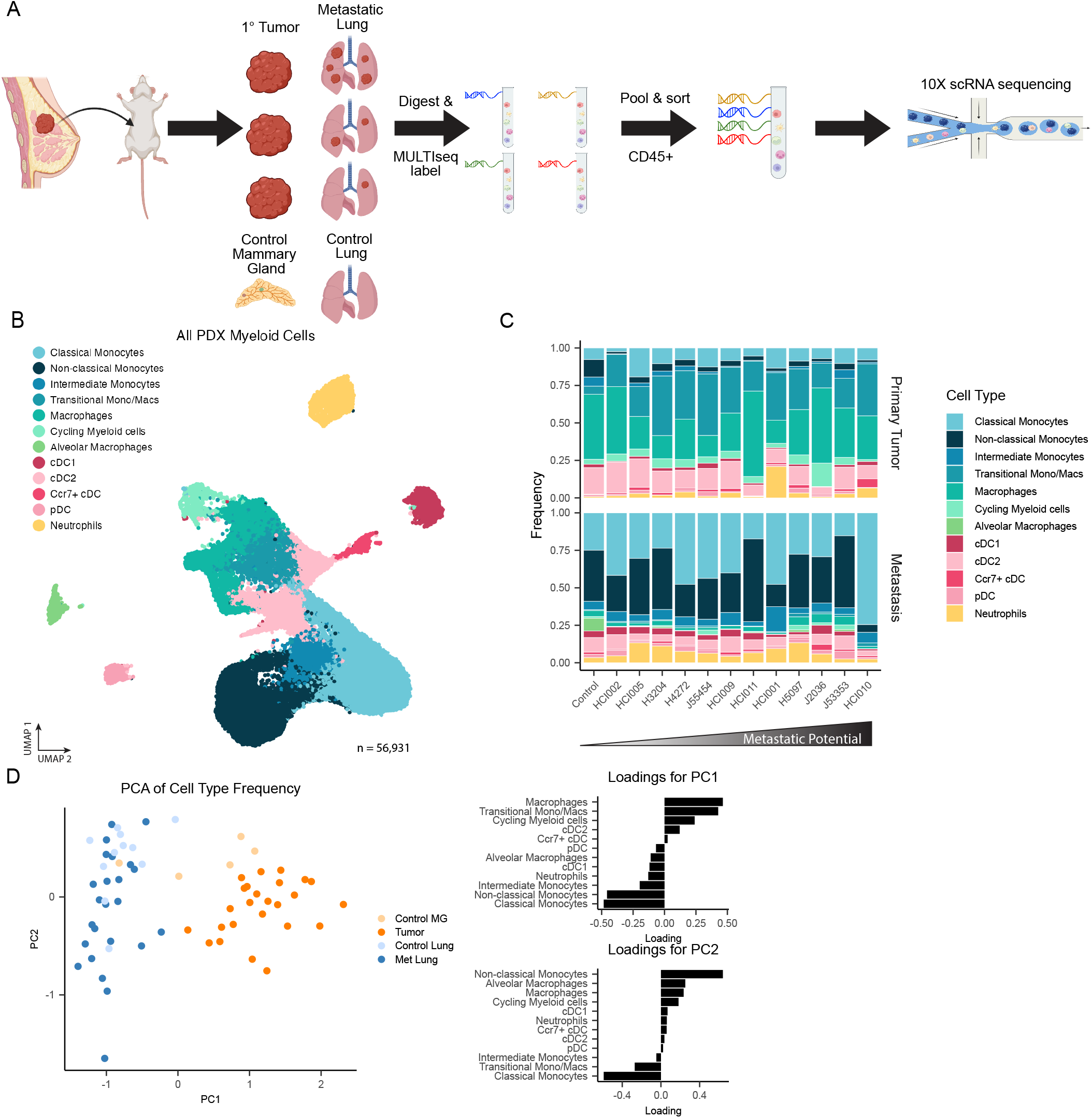
**(A)** scRNAseq experimental workflow. 12 Patient-derived xenograft models were orthotopically transplanted into the mammary gland of NSG mice. At 2.5cm primary tumor diameter tumors and metastatic lungs were harvested, along with control mammary glands and lungs from non-tumor bearing mice, and subject to digestion and labeled with MULTI-seq LMOs for multiplexing. CD45+ immune cells were sorted from all tissue samples, pooled, and sequenced using 10X Genomics scRNAseq platform. **(B)** UMAP of all major immune cell types in PDXs and control tissues sequenced. **(C)** Proportion of major immune cell types in each PDX model. For each PDX model and tissue type, cells from all samples sequenced were aggregated and proportion of each cell type was calculated as a fraction of all immune cells per PDX model and tissue. **(D)** Principal Component Analysis (PCA) of transformed cell type proportions per sample. First two PCs are plotted and samples are colored by tissue type (left). Loadings (contributions of each cell type to PCs) for PC1 and PC2 are displayed (right).

Following dimensionality reduction and unsupervised clustering, cell type annotation was performed using canonical markers [16–24] (Supp. Fig. S1B). As expected, we detected all major myeloid cell types including monocytes (classical, non-classical, and intermediate), macrophages (transitional mono/macs, tumor-associated, and lung alveolar), dendritic cells (cDC1, cDC2, Ccr7+, pDC), neutrophils, and a cluster of cycling myeloid cells (Fig. 1B). Lymphoid cells were lacking, which is consistent with the immunocompromised nature of PDX models (NSG mice) [25].

To understand the heterogeneity of myeloid cells across tumor models, we quantified the proportion of each major cell type in each tumor model and tissue in our dataset (Fig. 1C). Some models had particularly unique immune cell compositions, such as HCI010 metastatic lungs, which contained a large fraction of classical monocytes, and HCI001 primary tumors, which had the largest abundance of neutrophils. To identify coordinated changes in cell type composition across samples, we performed principal component analysis (PCA) on the cell type proportions in each sample. We observed strong differences in cell type composition between primary and metastatic tissues, as reflected by the separation of these tissues along PC1 (Fig. 1D, left). Primary tumors were predominantly infiltrated by macrophages and cycling myeloid cells, whereas metastatic lungs were associated with elevated proportions of monocytes and, as expected, tissue-resident alveolar macrophages (Fig. 1D, right). Control tissues similarly separated along PC1, suggesting that the difference in monocyte/macrophage proportion was a tissue-intrinsic difference and not specifically driven by malignancy.

### Characterization of the myeloid microenvironment of PDX primary tumors and metastases

To characterize phenotypic shifts of myeloid cells in the primary tumor and lung metastatic niche in greater detail, we next subclustered each major cell type (monocytes, macrophages, DCs, neutrophils) and annotated the sub-populations represented in each group.

Monocytes were separated into six different subsets: interferon-responsive (*Ifit3b, Ifit1, Ifit1bl1*), inflammatory (*Cxcl10, Il6*), intermediate (*H2-Ab1, H2-Eb1, H2-Aa*), myeloid-derived suppressor cell (MDSC)-like (*Lrg1, S100a8, Mmp8*) [24], non-classical (*Cd300e, Ace, Nr4a1*) [26] and stress-responsive (*Hilpda, Stab1, Sdc4*) [17] monocytes (Fig. 2A, Supp. Fig. S2A). Though the proportions of monocytes were generally more abundant in metastatic lungs compared to primary tumors, stress-responsive monocytes were specifically enriched in primary tumors (Fig. 2B, Supp. Fig. S2B).

**Figure 2.**
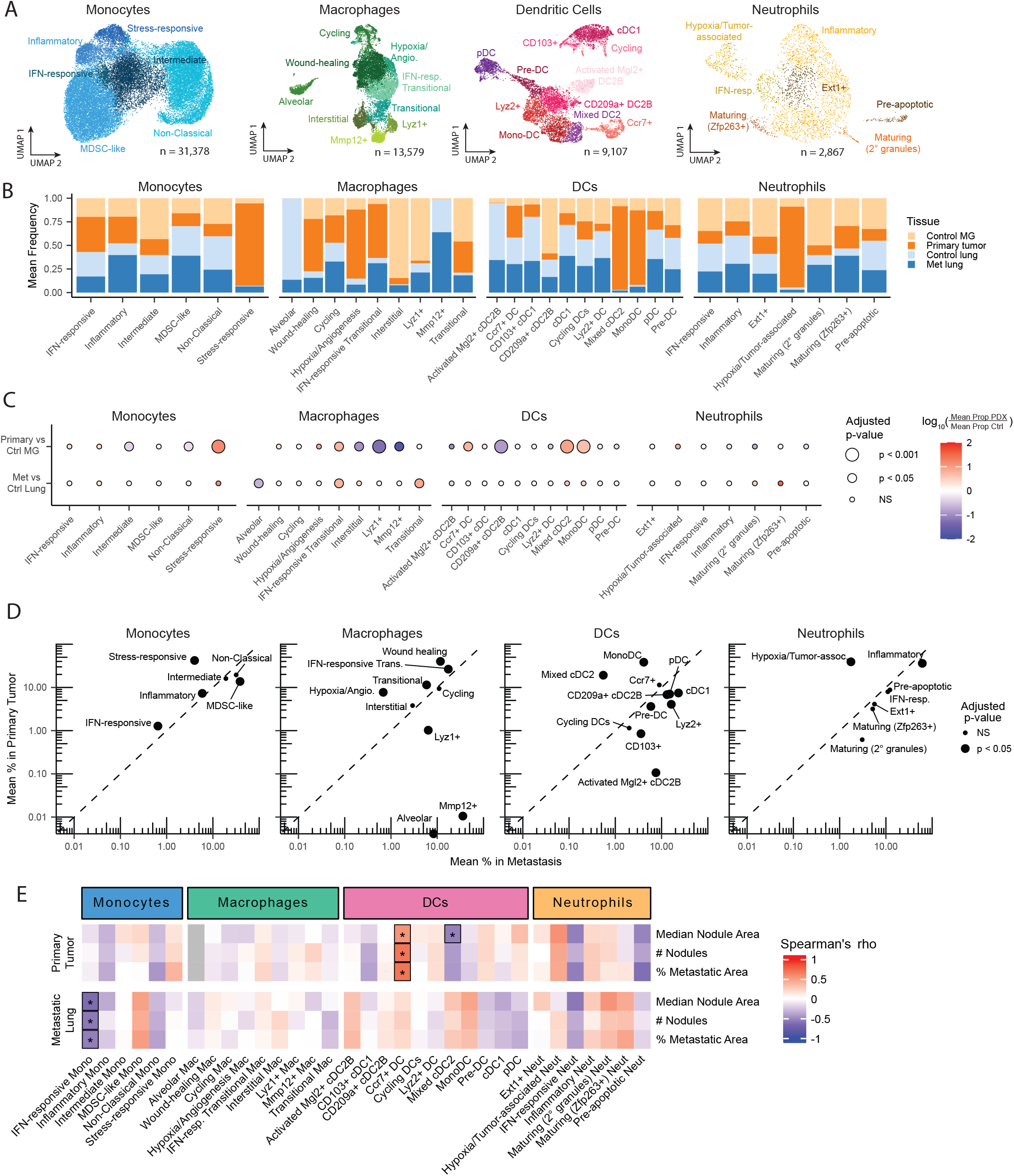
**(A)** UMAP of monocyte, macrophage, dendritic cell, and neutrophil sub-clustering, colored by subtype. **(B)** Tissue composition within each cell type subset. Objects for each major cell type were down-sampled to equalize total cell numbers across tissues prior to calculating proportions. Down-sampling was done 500 times and proportions were averaged across iterations. Mean CI widths (across all tissues and subsets) = 0.057 for monocytes, 0.031 for macrophages, 0.05 for DCs, 0.373 for neutrophils. **(C)** Log fold change of mean cell type subset proportion across malignant samples (metastasis or tumor) relative to corresponding controls. Dot size indicates significance of differential abundance between malignant and control tissues (linear mixed effects model with PDX as a random effect, followed by moderated t-test and Benjamini-Hochberg correction for multiple testing). **(D)** Average proportion of each cell type subset in metastatic lung vs primary tumor samples. Dot size indicates significance of differential abundance between primary and metastatic tumors (linear mixed effects model with PDX as a random effect, followed by moderated t-test and Benjamini-Hochberg correction for multiple testing). Dotted line indicates equal proportions. **(E)** Spearman correlation between cell type subset proportion and metastatic burden, split by tissue. Metastatic burden measures include median nodule area per sample, total number of nodules, and percentage of metastatic area as a fraction of total lung area (quantified by H&E). Significant correlations are indicated by * (Benjamini-Hochberg adjusted p-values).

Within the macrophage compartment we identified 9 subsets, many of which resembled previously described populations (Fig. 2A). These include hypoxic/angiogenic (*Egln3, Hilpda, Vegfa*) [17, 21], transitional (intermediate levels of *Ly6c* and *C1q*), IFN-responsive transitional (*Ifit1, Isg20, Ifit3b*, in addition to transitional markers) [17], cycling (*Ccna2, Ccnb1*), lung-resident alveolar (*Marco, Krt79*) [16], and Lyve1^+^ interstitial macrophages (*Lyve1, Folr2, Cd209f*) [22, 27, 28] (Supp. Fig. S2A). Another cluster highly expressing *Lyz1* and *Cd226* resembled an adipose tissue-associated macrophage found to be important in lipid homeostasis [29]. Accordingly, we found that these macrophages were primarily associated with control mammary gland tissues (Fig. 2B). Previously undescribed macrophage subsets were defined on the basis of their differentially-expressed marker genes (Supp. Fig. S2A). For example, the wound-healing cluster identified expressed genes such as *Hpgds, Col14a1*, and *Fgf13*. Lastly, we identified a group of macrophages highly expressing a number of ECM modulating genes (*Mmp12, Mmp13, P3h2, Adam22*), which was primarily composed of cells from control and metastatic lungs (Fig. 2B). Canonical tumor-associated macrophage (TAM) markers such as *Mrc1, Trem2, Arg1, Spp1*, were not specific to any one cluster (Supp. Fig. S2A).

Within the DC compartment, we identified 11 distinct subsets (Fig. 2A). This included two DC1 subsets distinguished by *Itgae* (CD103) expression; three DC2 subsets, including CD209a^+^ (*CD209a, Tnip3, Cd163*), activated Mgl2^+^ (*Ccl17, Adam23, Mgl2*), and mixed DC2A/B (*Ffar, Ltb*) [18]; as well as pre-DCs (*Tcf4, Cd7*) [30], monocyte-derived DCs (“monoDCs” — *Flt3, Fcgr1*), Lyz2^+^ DCs, and cycling DCs (Supp. Fig. S2A). Mixed DC2 and monoDCs were mostly isolated from primary tumors, whereas activated Mgl2^+^ DC2s and CD103^+^ cDC1s were enriched in metastatic and control lungs (Fig. 2B). Notably, monoDC enrichment in primary tumors relative to metastases has also been observed in human BC samples [31]. CD103^+^ cDC1s are a subset of tissue-resident DCs that migrate from peripheral tissues to lymph nodes to present antigen to T cells [32], and may represent airway pulmonary DCs that have been shown to play an important role in lung infections [33].

Within the neutrophil compartment, we identified 7 distinct subsets (Fig. 2A). The majority of cells were classified as inflammatory neutrophils (*Wfdc17, Adam19, Treml4*). Other clusters included IFN-responsive neutrophils (*Ifit3, Ifit3b, Ifi47*) [20], maturing neutrophils expressing secondary granule genes (*Camp, Ngf, Ltf*) [20], and hypoxia/tumor-associated neutrophils (*Mif, Hilpda, Ankrd37*) [16]. Hypoxic neutrophils were primarily isolated from primary tumors, while other neutrophil populations were similarly-associated with primary tumor and metastatic lung tissues (Fig. 2B). Three previously undescribed neutrophil subsets were defined on the basis of their marker genes (Supp. Fig. S2A). Ext1^+^ neutrophils expressed markers associated with vesicle trafficking and signaling at the plasma membrane (*Apba1, Cblb*). Another subset uniquely expressed *Zfp263*, which has been implicated in neutrophil maturation [34]. Lastly, the cluster of pre-apoptotic neutrophils expressed a number of genes related to NF*κ*B inhibition and apoptosis (*Nfkbid, Nfkbia, Dusp1*) [35].

Overall, these analyses demonstrate that PDX primary tumors and metastatic lungs consisted of both resident and infiltrating innate immune cells. These subpopulations were, in part, previously described in both homeostatic and diseased contexts. Furthermore, the observed tissue enrichment of myeloid subpopulations was largely in line with the reported functions of these cells (e.g. hypoxic myeloid populations in primary tumors, lipid macrophages in control mammary glands) [16, 17, 21, 29].

### Tissue-intrinsic differences dominate distinct myeloid phenotypes of primary and metastatic tumors

BC progression involves significant remodeling of myeloid cells. Therefore, we next calculated the fold-change in myeloid subtype proportions between malignant (tumor/metastasis) and control tissue samples (Fig. 2C). Generally, across all major cell types, we observed more consistent changes amongst primary tumors and control mammary glands compared to metastatic and control lungs (Supp. Fig. S2B, Supp. Fig. S2C). Accordingly, using the Bray-Curtis dissimilarity statistic [36] we found metastatic lung samples to be significantly more dissimilar to one another than primary tumor samples, particularly within the monocyte and macrophage compartments (Supp. Fig. S2D). This pronounced between-sample variability in metastatic lungs suggests that varying metastatic burden (a consequence of intertumor heterogeneity) leads to distinct remodeling of the metastatic niche. End-stage primary tumor tissues however, harbor a substantial, yet consistent, tumor burden across samples, which likely results in a more similar myeloid remodeling despite the intertumoral heterogeneity.

Among monocyte subsets, we observed a significant decrease in intermediate and non-classical monocytes in primary tumors, indicating either a differentiation shift or a decrease in trafficking of these cells during malignancy, and a significant increase in stress-responsive monocytes (Fig. 2C). Macrophage proportions were generally increased in malignant tissues compared to controls (transitional subsets most significantly), with the exception of alveolar macrophages [11] in the lung, and interstitial, Lyz1^+^, and Mmp12^+^ macrophages in primary tumors, which all decreased substantially compared to control tissues (Fig. 2C). DC abundance changes consisted of a significant depletion of CD209a^+^ DC2Bs, and enrichment of monoDCs, Ccr7^+^ DCs, and mixed DC2s in primary tumors relative to controls. This observation possibly reflects a general increase in the differentiation capacity of monocytes in mammary tissue, which mirrors the already described increase in macrophage populations. Finally, among neutrophil subsets, hypoxia-associated neutrophils had a trending increase in primary tumors relative to controls, consistent with the hypoxic microenvironment of primary tumors, whereas maturing (secondary granule) neutrophils had a trending decrease (Fig. 2C). However, no neutrophil subsets reached statistical significance, potentially due to the limited number of neutrophils detected in control samples.

To determine the unique myeloid cell composition of primary tumors and metastatic lungs, we compared the average proportion of each cell type subset between primary tumor and metastatic samples (Fig. 2D). In line with our prior analyses (Fig. 2B), primary tumors were enriched for stress/hypoxic subsets (monocytes, macrophages, and neutrophils), IFN-responsive subsets (monocytes and macrophages), and transitional/differentiating subsets (macrophages and DCs) relative to metastatic samples (Fig. 2D). The recurrence of hypoxic, IFN-responsive, and differentiation phenotypes within subsets of various myeloid cell types is indicative of a consistent microenvironment across primary tumors. Metastases, as noted before, showed higher inter-sample heterogeneity compared to primary tumors (Supp. Fig. S2D) and were enriched for MDSC-like monocytes, lung-specific macrophages (Mmp12^+^ and alveolar) (Fig. 2B), and the majority of identified DC subsets (Fig. 2D). Analogous comparison between control mammary gland and lung samples revealed that many changes were tissue-driven (Supp. Fig. S2E).

Despite the general tissue skewing, our analysis comparing compositional changes among control and malignant tissues revealed cell type subsets that were uniquely remodeled during malignancy. For example, Lyz1^+^ macrophages and CD209a^+^ DCs were depleted in control lungs compared to mammary glands. However, in malignant tissues these subsets were enriched in metastatic lungs compared to primary tumors (Supp. Fig. S2E). Conversely, Ccr7^+^ DCs were more abundant in control lungs compared to mammary glands, yet in malignancy they were enriched in primary tumors compared to metastatic lungs (Supp. Fig. S2E). Furthermore, this analysis also identified changes in subset proportions that, while trending in the same direction between control and malignant tissues, were significantly different in magnitude. For example, despite being more abundant in mammary tissues than lung tissues overall, interstitial and transitional macrophages were strongly depleted in primary tumors compared to control mammary glands (Supp. Fig. S2E). Thus, these cell type subsets represent distinct, stage-specific remodeling of the myeloid compartment, potentially mirroring the tumor cell-intrinsic changes accompanying metastatic progression [15, 37].

Our tumor models show varying metastatic potentials, which was determined by quantifying the number and size of metastatic nodules, and the total metastatic burden by histology [15]. These data provided the unique opportunity to identify myeloid subsets that are associated with different phases of metastatic progression (colonization vs outgrowth) [38, 39]. To this end, we analyzed the correlation between the abundance of individual myeloid cell subsets with the different measures of metastatic burden (Fig. 2E). Within primary tumor samples, the proportion of Ccr7^+^ DCs was significantly positively correlated with multiple measures of metastatic burden (median nodule size, total nodule number, and percent metastatic area, *ρ* = 0.5 to 0.7), while mixed cDC2 proportion was significantly negatively correlated with median nodule size (*ρ* = − 0.59). In metastatic lung samples, the proportion of IFN-responsive monocytes was negatively correlated with multiple measures of metastatic burden (*ρ* = − 0.56 to − 0.64), highlighting the already described anti-metastatic function of IFN-responsive monocytes [8]. Notably, this disease-associated shift in IFN-responsive monocytes was not captured by our tissue- and malignancy-level comparisons (Fig. 2C, Fig. 2D). Other cell types that had strong correlations with metastasis but did not reach significance include non-classical monocytes (*ρ*_Pct Met Area_ = − 0.39), MDSC-like monocytes (*ρ*_Pct Met Area_ = 0.5), and IFN-responsive neutrophils (*ρ*_Med Nod Area_ = − 0.51), confirming their known roles during metastatic progression [8, 10, 24].

In summary, the in-depth characterization of myeloid cell subsets and comparative analyses of BC PDX primary tumors and matched metastases identified cell types that were uniquely remodeled during metastatic progression (e.g. interstitial macrophages, Lyz1+ and CD209a+ DCs). Moreover, we identified cell subsets that positively (e.g. Ccr7^+^ DCs) and negatively (e.g. IFN-responsive monocytes) correlated with metastatic burden, suggesting pro- or anti-metastatic roles. Overall, these data demonstrate that many myeloid remodeling processes with known roles during primary tumorigenesis and metastatic progression were recapitulated in PDX models. Further, our approach additionally revealed many aspects of metastatic niche remodeling that was specific to PDX models with varying metastatic potential, providing insights into the role of tumor heterogeneity in the evolution of myeloid phenotypes during metastatic progression.

### Identifying metastasis-associated myeloid gene expression programs

Cell subtyping based on clustering restricts the discovery of functional cell states that may be associated with metastatic burden. Therefore, to characterize myeloid cell states in a cluster-independent manner, we applied DECIPHER-Seq [40] to each major cell type in our dataset (monocytes, macrophages, DCs, neutrophils) (Fig. 3A). DECIPHER-seq is an integrative non-negative matrix factorization (iNMF) approach, which identifies robust and unique gene expression programs and further harnesses inter-sample variability to identify modules (groups) of programs that are coordinated across samples [40]. Using DECIPHER-seq we identified 81 programs across all major cell types (25 monocyte programs, 25 macrophage programs, 23 DC programs, 8 neutrophil programs). These programs converged into three major modules of highly correlated programs from all major cell types, with module 1 more dominant in DC and monocyte programs, and module 2 more dominant in macrophage programs (Supp. Fig. S3A). Programs in module 1 and module 2 were largely enriched in lung/metastasis and mammary gland/primary tumor samples, respectively, confirming the already described tissue-specific myeloid cell composition (Supp. Fig. S3A). To assess if the modules converged on any common biological themes, we performed GSEA on all three modules using the Hallmark gene sets (Supp. Fig. S3B). Module 3 was significantly enriched for genes involved in mitosis. In line with this finding, DC Program 18 and Macrophage Program 10, which are part of module 3, were characterized by cell-cycle related genes (*Ccna2, Kif4, Ccnb1, Cdc20*). Module 2 was significantly enriched for several inflammation-associated gene sets, EMT, and hypoxia, which are all consistent with this module being enriched for primary tumor-associated programs. Module 1, however, was only weakly enriched for estrogen response genes and no other gene sets. We reasoned that the presence of fewer shared gene set enrichments for the lung-dominant module 1 might be due to the already described stronger inter-sample heterogeneity of metastatic lung tissues (as a result of the varying metastatic burden) than end stage tumors (Supp. Fig. S2D). Collectively, these data highlight tissue-specific coordination of myeloid phenotypes in primary tumors and metastasis.

**Figure 3.**
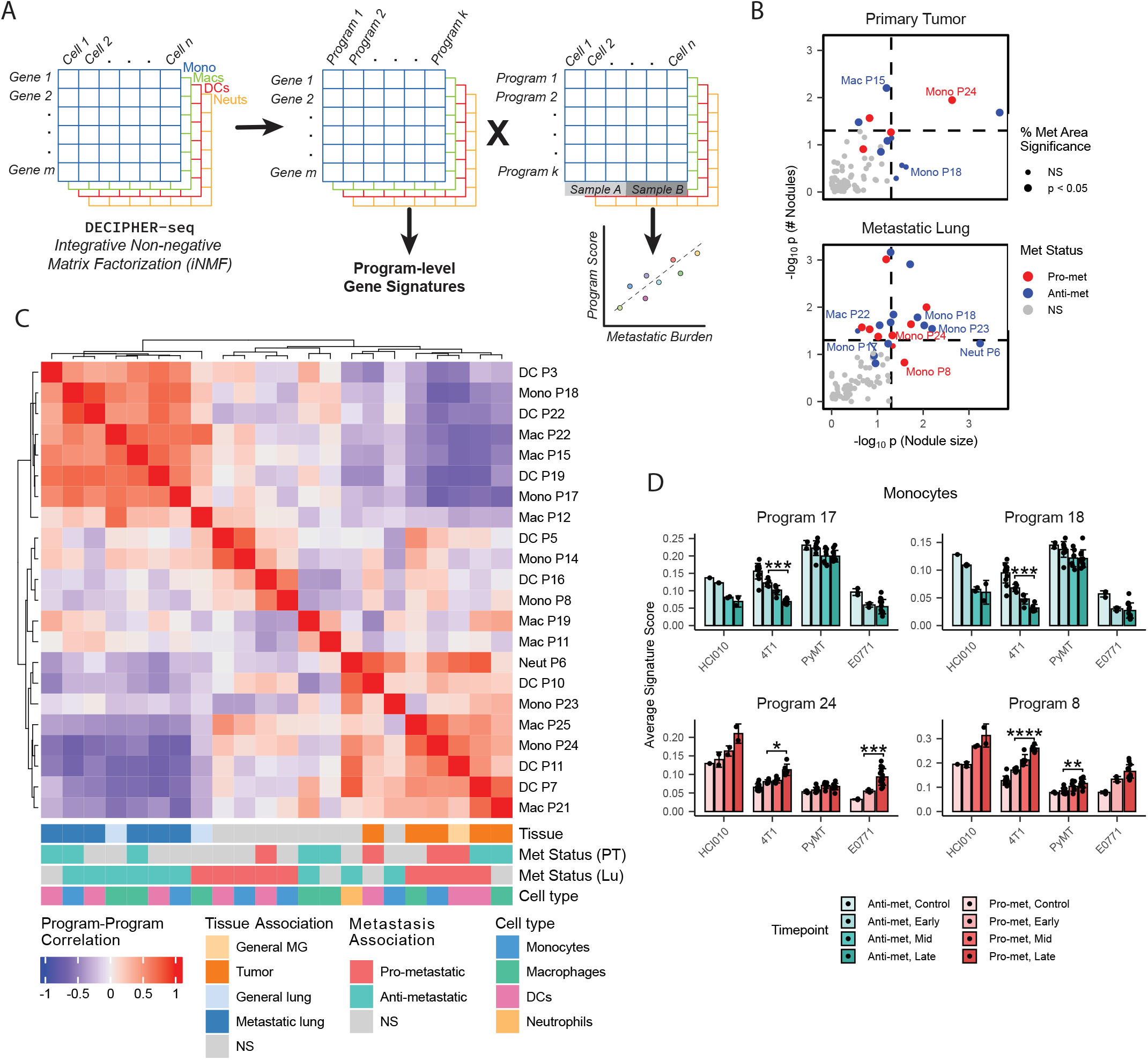
**(A)** Non-negative matrix factorization (NMF) analysis workflow. DECIPHER-seq was applied to all PDX immune cells, separately for each major cell type (monocytes, macrophages, DCs, neutrophils). Gene by program matrices were then used to create gene signatures, using the top 50 marker genes, for each NMF program. Program by cell matrices were used to perform statistical testing between program scores and different metadata features. **(B)** -log10(p-value) of the correlations between NMF program score and either median nodule size or total number of nodules across all samples. Point size indicates whether programs were additionally significantly correlated with percent metastatic area. Correlations were performed separately for tumor and metastatic samples. Programs are colored by positive or negative correlation across all metastatic burden measurements. **(C)** Pairwise correlation between all metastasis-associated NMF programs. Programs are annotated according to their tissue status, metastasis status (per tissue), and cell type. **(D)** Average signature score of selected monocyte NMF programs in four publicly available time course datasets. Signature scores were calculated for monocytes in each sample, and samples were grouped according to time point. Time-points were as defined in original publications.

Next, we performed statistical analyses to identify programs that were differentially associated with the different metadata features of our dataset, such as cell cycle score and metastatic burden (i.e. number and size of nodules, percent metastatic area) (Fig. 3A). To test the robustness of our strategy, we first focused on DC Program 18 and Macrophage Program 10, which were part of proliferation module 3. Both programs were significantly correlated with cell cycle score, motivating their characterization as “proliferation programs” (Supp. Fig. S3C). With our workflow successfully benchmarked, we next identified several gene expression programs across cell types that were significantly positively or negatively correlated with metastatic burden, and therefore categorized as pro- or anti-metastatic programs (Fig. 3B). For example, neutrophil Program 6 and monocyte program 23 were negatively correlated with metastasis and expressed a large number of IFN response genes (Fig. 3B, Supp. Fig. S3D), reminiscent of a previously described anti-metastatic IFN-driven mechanism of tumor model HCI002 [8]. Besides HCI002, a number of other lowly metastatic models, such as H4272 and H3204, also expressed these programs, thus extending this anti-metastatic phenotype to other BC models in a more general manner (Supp. Fig. S3E).

While carefully evaluating the correlations of the gene expression programs with the different histological measures of metastasis (number of metastatic nodules, nodule size, metastatic burden), we found that several metastasis-associated programs were exclusively significantly correlated with either median nodule size or total nodule number. For example, monocyte program 8 contained several marker genes commonly expressed by MDSCs and was significantly positively correlated with nodule size (*ρ* = 0.45) but not nodule number (*ρ* = 0.30) (Supp. Fig. S3D, Fig. 3B). Conversely, macrophage program 22 contained several marker genes characteristic of alveolar macrophages and was significantly negatively correlated with nodule number (*ρ* = − 0.52) but not nodule size (*ρ* = − 0.28) (Supp. Fig. S3D, Fig. 3B). Individual nodules are thought to represent distinct seeding events [38, 39]. Therefore, this finding is particularly intriguing because it suggests that different myeloid cell phenotypes may play distinct roles in either colonization/seeding or outgrowth, and that these two “phases” of metastasis may be targetable in different ways.

### Evolutionary switch from anti-to pro-metastatic myeloid gene expression programs during metastatic progression

To further understand how the metastasis-associated programs are coordinated across samples, we applied unsupervised clustering to their DECIPHER-seq pairwise correlation matrix (Fig. 3C). Programs clustered according to their tissue association, however, we were surprised to find that tissue and metastasis association were concordant. Most primary tumor-associated programs were pro-metastatic, whereas most lung-associated programs were anti-metastatic. The tissue-metastasis concordance was unlikely an artifact of calculating correlations for each tissue separately, since both pro- and anti-metastatic programs appeared in both tissue types (Fig. 3C, “Met Status” annotation bars). Additionally, programs from all cell types were comparably represented within both lung/anti-metastatic and primary tumor/pro-metastatic modules, regardless of the already observed tissue skewing of cell type abundances (Fig. 3C, “Cell type” annotation bar).

We therefore began to search for other explanations as to why tissue and metastasis associations might be concordant. One variable that distinguished both tissue types and metastasis association categories was tumor burden. By definition, pro-metastatic programs were correlated with higher metastatic burden, and anti-metastatic programs were correlated with lower metastatic burden. Similarly, primary tumors reflect a much higher tumor burden than metastatic lungs, even for the most highly metastatic samples. Therefore, we hypothesized that these myeloid gene expression programs reflected tumor burden and were potentially the result of a local remodeling by tumor cells. Consequently, we would expect these gene expression programs to change over time with increasing metastatic burden. To validate the presence of our identified metastatic programs and evaluate their temporal dynamics during metastatic progression, we created gene signatures for each program and scored publicly available time-course datasets of metastatic lung myeloid cells isolated from highly metastatic PDX (HCI010), and immune-competent orthotopically transplanted (4T1, E0771) and genetically engineered (PyMT) mouse models of BC [11–13]. Metastasis-associated monocyte programs were most frequently significantly altered over time (Supp. Fig. S3F). Most DC programs did not validate in the syngeneic datasets, perhaps affected by the lack of T cells in PDX models. Additionally, the PyMT and E0771 models generally showed weaker expression of many gene expression programs compared to the 4T1 and HCI010 datasets. Of note, PyMT and E0771 are classified as luminal BC subtype [41,42], while HCI010, 4T1 [43], and the majority of the models in our dataset [15, 44] are triple negative. As previously reported [11], many pro-metastatic monocyte programs significantly increased over time in the metastatic lung (Fig. 3D). In contrast, anti-metastatic monocyte programs were present in the pre-metastatic niche and decreased with metastatic progression. This suggests that even in highly metastatic tumor models, the immune system may initially repress metastasis.

In summary, by leveraging histological variables [15] in these heterogeneous PDX models along with NMF analysis, we identified both known and novel myeloid gene expression programs that were associated with metastatic phenotypes and dynamically changed during metastatic progression. We find that these programs are coordinated on a tissue level and likely represent distinct roles during metastatic seeding and outgrowth. Moreover, many of our identified metastasis-associated programs validated in immunocompetent metastatic mouse models of BC, and pro- and anti-metastatic programs showed inverse temporal dynamics representing an evolutionary switch in myeloid phenotypes during metastatic progression.

### Distinct monocytic MDSC phenotypes reside in primary tumors and metastatic lungs

Of the pro-metastatic programs, monocyte programs 8 and 24 most significantly increased over time and were strongly correlated with metastatic burden (Supp. Fig. S3F, Supp. Fig. S4A). Programs 8 and 24 were highly concordant and correlated with a monocytic MDSC signature (Fig. 4A, Fig. 4B, Supp. Fig. S4B), yet they were expressed in largely distinct sets of cells (Fig. 4A, Supp. Fig. S4C). Furthermore, on the sample-level, program 8 was more active in metastatic lungs, whereas program 24 was more active in primary tumors (Fig. 4C, Supp. Fig. S4D). Monocytic MDSCs (M-MDSCs) can have various roles in promoting metastasis [5, 45]. However, it is unclear whether these different roles of M-MDSCs are linked to underlying transcriptional heterogeneity. Therefore, we hypothesized that M-MDSC-linked monocyte programs 8 and 24 may have distinct roles and phenotypes during metastasis.

**Figure 4.**
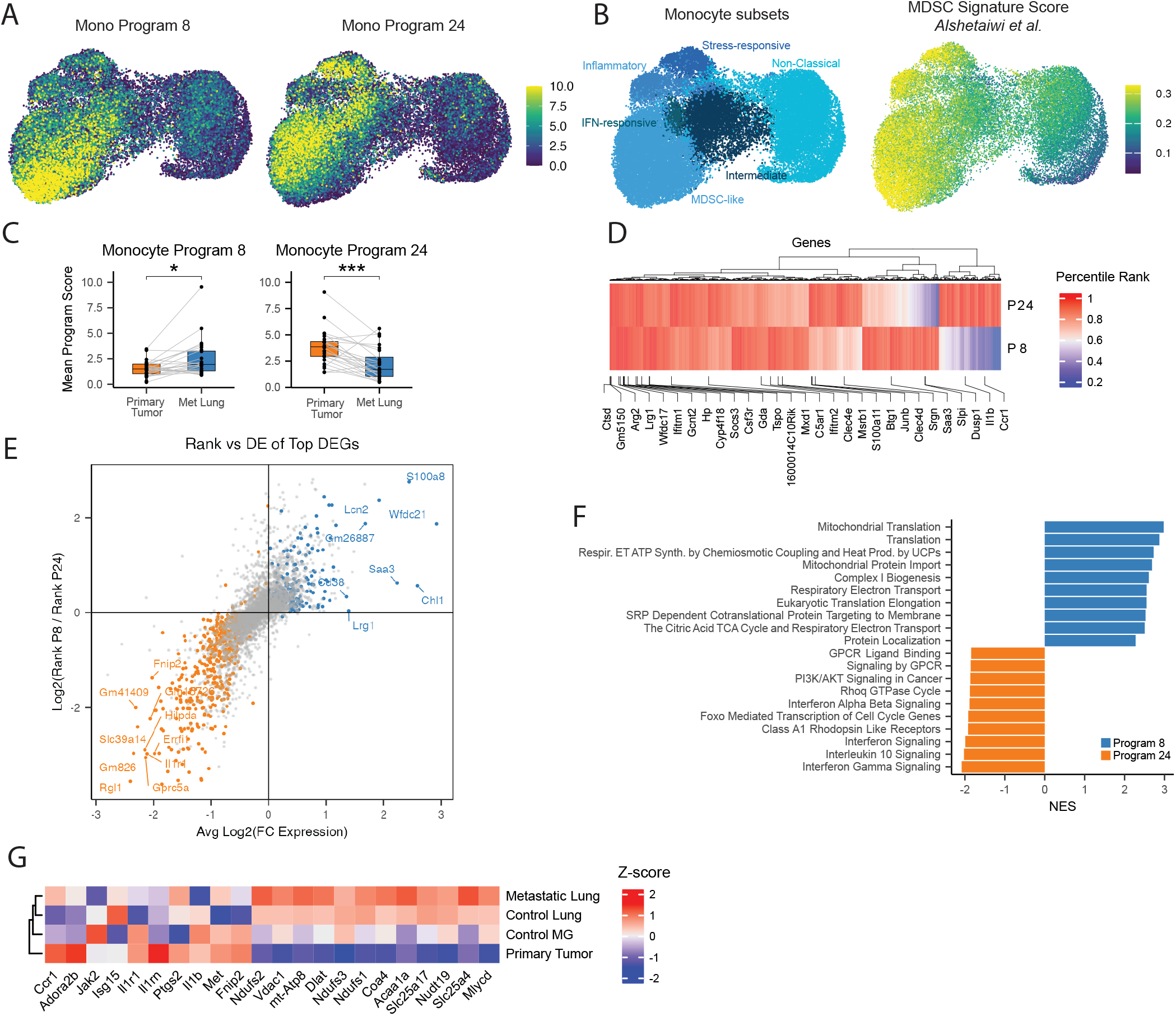
**(A)** UMAP of program 8 (left) and 24 (right) score across all monocytes. **(B)** UMAP of monocyte subsets (left), and UMAP of M-MDSC Signature score from Alshetaiwi et al. (right). **(C)** Comparison of metastatic lung and tumor average program score per sample for monocyte programs 8 and 24. T-test for Program 8, Wilcox test for Program 24, * = p < 0.05, *** = p < 0.001. **(D)** Percentile rank of top 10% of genes (by NMF weight) in programs 8 and 24 (n = 1,687). Annotated are M-MDSC signature genes (from Alshetaiwi et al.). **(E)** Comparison of average log2 fold-change gene expression and gene ranking by NMF weight. Annotated genes are among the top fold-change for each program. Color indicates significance from differential expression testing. **(F)** Gene set enrichment analysis of differentially expressed genes between program 8 and 24 top activity cells using Reactome gene sets. Top 10 significant (padj < 0.05) pathways are shown sorted by NES. **(G)** Z-scored expression of selected genes from top pathways for program 8 and 24 amongst monocytes, pseudobulked by tissue.

To further interrogate the differences between monocyte programs 8 and 24, we compared the rankings (by NMF weight) of genes in program 8 and 24 followed by unsupervised clustering (Fig. 4D). This analysis resulted in three main clusters of genes: one group consisted of genes that were highly ranked in both programs and included many known MDSC signature genes, and two other groups consisted of program-specific genes. However, gene ranking by NMF weight is not necessarily indicative of gene expression [46, 47]. Therefore, to identify expression-level differences between programs 8 and 24 we performed differential gene expression (DGE) analysis between cells whose activity was most driven by these programs (Supp. Fig. S4E, Supp. Fig. S4F). Accordingly, the genes identified using DGE were generally highly ranked in their respective programs, but they were not necessarily the highest-ranked genes (Fig. 4E). We next performed GSEA on the DEGs between program 8 and 24. Program 8 MDSCs upregulated genes related to oxidative phosphorylation, whereas Program 24 MDSCs upregulated genes related to IFN response, and chemokine production/signaling (Fig. 4F). In macrophages, OXPHOS supports wound-healing/anti-inflammatory functions, while glycolysis supports pro-inflammatory functions such as *Il1b* secretion [48, 49]. In line with this finding, we observed that several genes, such as *Il1b, Ptgs2* (COX2), *Il1r1*, and components of the electron transport chain (e.g. *Ndufs3, Ndufs1*), were differentially expressed by monocytes in metastatic lungs and primary tumors (Fig. 4G).

Our data revealed two distinct pro-metastatic M-MDSC subsets that were either enriched in primary tumors or metastases. M-MDSCs in primary tumors increased inflammatory signaling and metastasis-enriched M-MDSCs showed altered metabolism and increased oxidative phosphorylation. Both populations correlated with metastatic burden and dynamically increased during metastasis progression.

### Decrease in classical to non-classical monocyte transcriptional programs coincides with metastatic progression

Of the anti-metastatic monocyte programs, Programs 17 and 18 were most significantly decreased over time (Fig. 3D), so we chose to characterize them more indepth. Both programs were significantly negatively correlated with percent metastatic area (Fig. 5A), and highly correlated between each other across samples (Fig. 5B), suggesting they were coordinated in their anti-metastatic phenotype. Program 17 was expressed across several monocyte subsets, primarily non-classical, intermediate, and inflammatory monocytes, whereas program 18 was primarily expressed by non-classical monocytes (Fig. 5C). Non-classical monocytes have been shown to play an anti-metastatic role by recruiting and activating NK cells [10]. Notably, NK cells were lacking in our models, suggesting additional anti-metastatic functions of non-classical monocytes. Further, nearly all samples had some distribution of classical, intermediate and non-classical monocytes and program 17 and 18 expression (Supp. Fig. S5A, Supp. Fig. S5B), indicating that these programs reflect a more general anti-metastatic remodeling independent of tumor heterogeneity.

**Figure 5.**
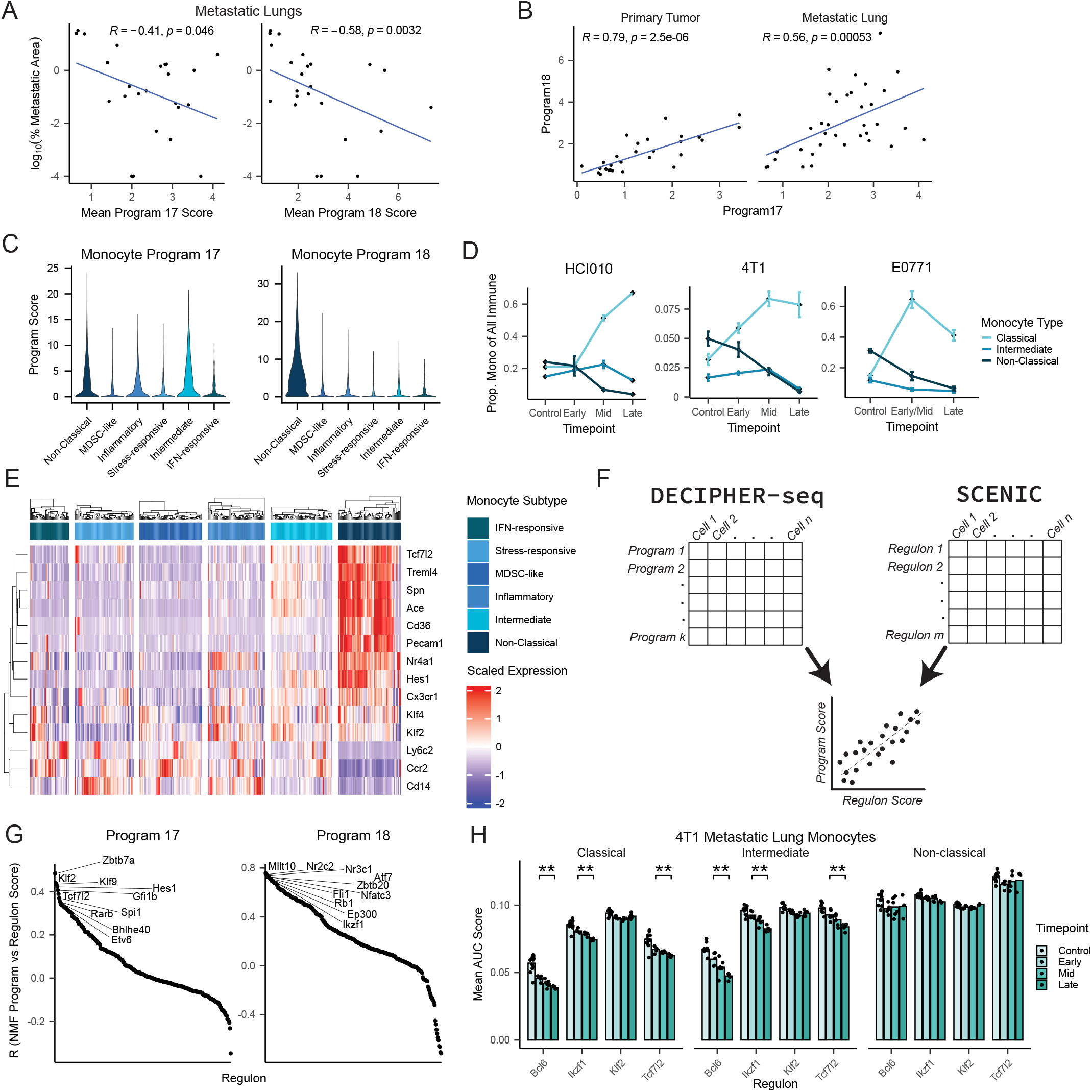
**(A)** Spearman correlation between average program 17 and 18 score and percent metastatic area in the lungs. Program scores were averaged across all monocytes within each sample. Only metastatic lung samples are shown. **(B)** Spearman correlation between average program 17 and 18 score (across all monocytes) per sample. Correlations are split according to tissue. **(C)** Distribution of program 17 and program 18 scores across all monocyte subsets. **(D)** Proportion of major monocyte subsets (classical, intermediate, non-classical) in the lungs of metastatic breast cancer models (HCI010, 4T1, E0771) at different timepoints. Monocyte proportion was calculated as a fraction of all immune cells. Mean and standard error for each timepoint are shown. **(E)** Expression of classical (Cd14, Ccr2, Ly6c2) and non-classical monocyte markers across various PDX monocyte subsets. Expression pseudobulked by sample and monocyte subset. **(F)** Strategy for determining transcriptional regulators of NMF programs. Correlation between DECIPHER-seq program scores and SCENIC regulon activity scores were calculated across all cells to identify candidate transcriptional regulators of gene expression programs. **(G)** Correlation between each regulon and program 17 (left) and 18 (right) across all monocytes. Regulons with the highest correlations are annotated, in order. **(H)** Average regulon activity scores (mean AUC) for selected transcription factors associated with classical to non-classical monocyte differentiation. Average regulon activity scores were calculated across all classical, intermediate, and non-classical monocytes per sample. Bars represent average scores across all samples for each timepoint. Wilcoxon rank-sum test was used to test for significance, Benjamini Hochberg correction was applied to calculate adjusted p-values (* p < 0.05, ** p < 0.01)

Interestingly, time-course experiments [11–13] of lung metastatic progression in BC consistently showed decreased abundance of non-classical monocytes, and to a lesser degree intermediate monocytes, while classical monocytes increased over time (Fig. 5D). Non-classical and intermediate monocytes differentiate from classical monocytes [50]. Though we cannot exclude the possibility that fewer intermediate or non-classical monocytes were trafficking to the lung, we hypothesized that this program represented a differentiation-associated program given the activity of program 17 spanned the monocyte differentiation trajectory.

Program 17 was characterized by genes that are typically associated with non-classical monocytes, but were expressed to varying degrees across multiple classical monocyte subsets, primarily in inflammatory monocytes (Fig. 5E). To further investigate if program 17 was involved in monocyte differentiation, we performed SCENIC gene regulatory network analysis [51]. To identify potential upstream regulators of gene expression programs, we calculated the correlation coefficient between each transcription factor (TF) regulon’s score and gene program scores (Fig. 5F). As a proof of concept, IFN programs (monocyte program 23 and neutrophil program 6) were significantly and highly correlated with known drivers of IFN signaling, such as *Stat1, Stat2*, and *Irf9* [52] (Supp. Fig. S5C). Likewise, cell-cycle associated programs (macrophage program 10 and DC program 18) were significantly and highly correlated with known transcriptional drivers of cell cycle progression, such as *Mxd3 (Mad3), E2f8*, and *E2f7* [53, 54] *(Supp. Fig. S5C)*.

Many of the top significantly correlated TFs for program 17 were TFs previously described to be important for classical to non-classical monocyte differentiation, such as *Klf2, Tcf7l2*, and *Bcl6* [26, 55, 56] (*Fig. 5G)*. We also identified several TFs as upstream regulators of program 17 that have not previously been implicated in classical to non-classical monocyte differentiation, such as *Zbtb7a* and *Gfi1b*, suggesting their involvement in the pro-metastatic differentiation shift of monocytes. We independently performed SCENIC on monocytes from a 4T1 time-course dataset of the metastatic lung and we found many of these TFs decrease significantly over time (Fig. 5H, Supp. Fig. S5D). Notably, the decrease in TF activity was only seen in classical and intermediate monocytes, and not in non-classical monocytes, suggesting that the decrease in TF activity was not driven by decreases in absolute number of non-classical monocytes after differentiating (e.g. apoptosis), but rather changes in precursor cells. Collectively, these data provide evidence of a yet undiscovered shift in monocyte differentiation-associated transcriptional programs within the lung that coincide with metastatic progression.

## Discussion

In the present study, we characterized the heterogeneity of myeloid cells across breast primary tumors and their matched metastatic lung tissues within PDX models with varying metastatic phenotypes. This comparative single-cell analysis revealed that primary tumors and metastatic lungs have marked differences in their myeloid cell compositions, even when accounting for baseline differences in immune composition (macrophage- vs monocyte-dominated, respectively). Differences in myeloid subset proportions were largely tissue-driven, but also included malignant-specific remodeling that was either specific to primary tumors or metastases.

Our findings are in line with prior work comparing the primary and metastatic TME in the E0771 BC model, which also identified shifts in immune cell composition between primary tumors and lung metastases [13]. However, the specific myeloid cell types that change between primary and metastatic E0771 tumors were largely different from those seen in our heterogeneous PDX models. For example, in E0771 tumors IFN-responsive macrophages were more abundant in metastases, and intermediate monocytes and migratory DCs were more abundant in primary tumors. Conversely in our data, IFN-responsive macrophages were most abundant in primary tumors, and intermediate monocytes and migratory DCs were not significantly different between the two tissues. It’s possible that the difference in mouse strain, immunocompetence, or increased heterogeneity in PDX models compared to established cell lines like E0771, may play a role here. Additionally, our dataset recapitulates several malignancy-induced changes which have previously been described in other human and murine studies, such as an enrichment of hypoxia-associated myeloid cells in primary tumors and depletion of tissue-resident macrophages in both primary and metastatic sites [11, 17, 21, 57, 58]. Importantly, in contrast to previous work, we carefully disentangled effects of malignancy-induced changes vs tissue-intrinsic differences [13, 31]. Our results suggest that tissue, and not tumor progression, is the dominant driver of alterations to immune cell composition between the primary tumor and the metastatic site.

We subsequently identified gene expression programs that were associated with metastatic progression. Though myeloid cells have diverse and sometimes paradoxical roles in primary tumorigenesis and metastasis, these features can be better resolved when placed in the context of specific metastatic phenotypes. Crucial to identifying these metastasis-associated programs was the detailed histological characterization of the number of nodules, size of nodules, and total metastatic area in the lung, which revealed that certain gene expression programs were specifically correlated with the number or size of metastatic nodules. Examples of such programs include an alveolar macrophage program negatively correlated with the number of nodules, and an MDSC-like program positively correlated with the median size of nodules. These results are in line with recent studies showing alveolar macrophages play important roles early in lung colonization [57], and that MDSCs can secrete many growth promoting factors that help tumor cells successfully grow in the metastatic site [5, 45]. Importantly, this result emphasizes the importance of how metastasis is quantified. Our results implicate that these metrics may represent distinct biological processes, and that myeloid cells may play different roles in seeding/early stages of disseminated tumor cells (as measured by number of metastatic nodules), and outgrowth of metastatic colonies in the lung (as measured by size of metastatic nodules).

Using time-course data, we validated pro- and anti-metastatic programs in immune competent, syngeneic, and genetically modified mouse models (4T1, E0771, PyMT) and found that these programs dynamically changed during metastatic progression. It is well known that primary tumors remodel distant tissues to favor metastatic colonization [59]. Accordingly, we found two M-MDSC programs positively correlated with metastasis, one more active in primary tumor monocytes and the other in metastatic lung monocytes. Interestingly, though these two gene programs shared expression of many core M-MDSC genes, they were expressed in distinct groups of cells, and differentially expressed a subset of genes relating to pro-inflammatory signaling and mitochondrial metabolism. A recent single-cell profiling of human glioblastoma (GBM) identified two M-MDSC states that are strikingly similar to the ones we identify here [60]. In this study, IDH wildtype GBMs have low rates of metastasis, but are particularly invasive to surrounding brain tissue, and the authors suggest these distinct MDSC states may be associated with GBM invasiveness. Our work not only extends these phenotypes to metastatic BC models, but also supports the nascent notion that there is both functional and phenotypic heterogeneity within M-MDSCs.

Conversely, the dynamics of anti-metastatic cell types within the pre/early-metastatic niche are currently not very well understood. The idea that pro- and anti-metastatic cell types might co-exist within the pre-metastatic niche is not entirely unprecedented [12]. A previous study temporally profiling 4T1 metastasis to the lung also found a decrease in type I IFN response in monocytes over the course of metastatic progression [14], similar to the one we identified in our dataset and found to be decreased over time in HCI010 and 4T1 models [11, 12].

We also propose a novel mechanism explaining why anti-metastatic non-classical monocytes decrease over time. Non-classical monocytes have been shown to be important in suppression of metastasis via NK-mediated killing of tumor cells in the vasculature [10]. However, with the absence of NK cells in NSG mice we were surprised to find non-classical monocyte programs negatively correlated with metastasis. We show both in immunocompetent and immunodeficient models that non-classical monocytes decrease in abundance, and that TFs known to regulate non-classical monocyte differentiation (e.g. *Tcf7l2, Klf2, Bcl6*) [26, 55, 56, 61] decrease over time in the metastatic niche, specifically within classical and intermediate monocytes. These data suggest a stalled monocyte differentiation caused by decreased TF activity, resulting in a compositional shift from non-classical to classical monocytes during metastatic progression. We acknowledge several limitations of this analysis, including not knowing the relative contributions from non-classical monocytes that differentiate locally in the lung versus those that differentiate in the bone marrow and transit to the lung. However, given emerging evidence on myeloid differentiation biases during tumorigenesis [62], our data provide compelling evidence to motivate future studies on monocyte differentiation skewing during metastasis and how it may be therapeutically targeted.

In summary, our findings highlight the complexity and heterogeneity of myeloid remodeling during metastatic progression, and uncover a progressive shift from anti-to pro-metastatic myeloid phenotypes within the metastatic lung. Future work will focus on understanding how these myeloid programs interact with the native microenvironment, such as cells of the adaptive immune system, and functionally validate the metastasis-associated phenotypes identified here. Our study provides novel insights into how to develop therapies preventing metastases in high-risk patients by harnessing the anti-metastatic remodeling present during pre-metastatic niche development and blocking their transformation into pro-metastatic phenotypes.

## Methods

### Animal studies

PDX primary tumor and metastatic lung tissues were generated from novel and established PDX models as previously described [15]. The UCSF Institutional Animal Care and Use Committee (IACUC) reviewed and approved all animal experiments. Patient-related information of PDX models were described previously [15]. Briefly, generation and propagation of PDX models was done by orthotopically transplanting 1-mm-thick PDX tumor pieces into cleared mammary fat pads of 4-week-old NOD-SCID gamma mice (NSG), as previously described [44, 63, 64]. Animals were euthanized when the primary tumor reached 2.5 cm in diameter. At endpoint, metastatic lungs were perfused and primary tumors and lungs were collected, cut into small pieces, and cryopreserved using Recovery Cell Culture Freezing Medium (Thermo Fisher #12648010). Cryopreserved samples were stored in liquid nitrogen until preparation for sequencing.

### Sample Preparation

All tumor and lung samples were processed as previously described [15]. For control mammary glands, viable tissues were thawed from frozen and briefly washed in a filter with DMEM/F12 media. Mammary glands were then minced with a scalpel and dissociated in a digestion mix as described [15] using a standard GentleMACS protocol (37_m_TDK1). After digestion, samples were spun at 300 × g for 10 mins at 25°C. After centrifugation, all pipettes were precoated with a BSA solution prior to contacting cell suspensions. The top (fatty) layer was removed, transferred to a separate tube with DMEM/F12, and agitated to disperse the fat before re-spinning (as before) to collect any remaining cells. The pellets from both spins were pooled and gently mixed with DNase (2000 U/mL) for 3–5 minutes before spinning again (as before). Cells were resuspended in PBS and counted prior to FACS staining and MULTI-seq labeling (as previously described [15]).

### Batch Design

This dataset was sequenced across six batches (PDX1, PDX2, PDX3, PDX7, PDX9, PDX12). Library preparation and sequencing slightly differ between batches. Namely, PDX1-3 library preparations were performed using the 10X v2 library kit (details can be found in [15]), whereas PDX7, 9, and 12 were performed as described below. Raw feature and MULTI-seq barcode FASTQs for all batches were uniformly pre-processed and demultiplexed for this study (further details below) to ensure consistency across the dataset.

### Library Preparation and Sequencing

Library preparation was performed using the 10X v3 library kit with modifications to generate MULTI-seq libraries, as previously described [12]. Libraries were sequenced on a NovaSeq 6000 or NovaSeqX Sequencing System (Illumina) using standard protocols for 10X libraries.

### scRNAseq Data Pre-processing and QC

FASTQs were pre-processed using Cell Ranger count default parameters (v7.1.0, 10X Genomics) and aligned to the combined human and mouse GRCh38 and mm10 2020-A reference transcriptome. “Species-aware QC” involved filtering feature-barcode matrices for cells with *>*90% mouse reads, and mouse genes with *>*3 total UMI counts present in *>*3 cells. General quality control using Seurat (v5.3.0) included additionally filtering for cells with *<*5% mitochondrial reads, *>*800 total counts and *>*250 genes.

### Sample Demultiplexing

Sample demultiplexing was performed using deMULTIplex2 (v1.0.2) [65]. For batches PDX1, 2, and 3 which were sequenced using the 10X v2 kit, start and end indices of cell barcode, sample tag, and UMI sequences differed from the default and were therefore specified in the readTags function (cell barcode: 1–16; sample tag: 1–8; UMI: 17–26). For all batches, read tables were filtered for cells in the MULTI-seq FASTQ filtered feature barcode matrix. Tag matrices constructed from the read tables were then filtered for cells that passed QC as described above. Demultiplexing (using the ‘demultiplexTags’ function) was performed using the default parameters. The following dataset-specific adjustments were made to maximize cell assignment:

- PDX1 and PDX3 batches were demultiplexed as previously described [15].
- For batch PDX12, barcode libraries were sequenced across multiple NovaSeq lanes so readtables were concatenated across sequencing lanes prior to creating tag matrices.

After running deMULTIplex2, samples with fewer than 25 assigned “singlets” per batch were removed (which applied to two samples pertaining to the PDX1 batch). An additional doublet filtering step was performed whereby cells assigned as “multiplet” or “singlet” from all batches were combined in a Seurat object and subset by major cell type. Cell type subset objects were then clustered, and clusters containing ∼ *>*60% “MULTI-seq defined multiplets” were removed, as previously described [11]. Furthermore, at this stage we also removed any non-immune cells detected by marker analysis. The resulting quality-controlled, de-multiplexed and doublet-removed data was used for integration and cell type annotation.

### Data Integration and Clustering

After removal of multiplets and unassigned cells, the full Seurat object was split by batch (“experiment” in the object metadata) and processed in the following manner: Raw counts were log normalized with a scale factor of 10,000 (`NormalizeData` function). Top 2000 variable features were identified using the VST method (`FindVariableFeatures` function). All genes were then scaled to a maximum value of 10 (`ScaleData` function). Principal component analysis was performed to calculate the first 50 principal components (`RunPCA` function).

To determine the number of PCs that captured the majority of the variability in the dataset we took a quantitative approach as described by the Harvard Chan Bioinformatics Core (https://hbctraining.github.io/scRNA-seq/lessons/elbow_plot_metric.html). We selected the minimum of two values: either the number of PCs which cumulatively contribute greater than 90% of the standard deviation and individually contribute to less than 5% of the standard deviation, or the PC where the percent change in standard deviation is less than 0.1%.

We then performed Harmony integration using the `IntegrateLayers` function. Layers were then joined using the `JoinLayers` function and the number of harmony components were selected (as described for PCs). Subsequently, we performed UMAP dimensionality reduction and determined the k-nearest neighbors with the selected number of harmony components (`RunUMAP`, `FindNeighbors` functions). Clustering was performed using the leiden algorithm and the igraph method across a range of resolutions (typically ∼ 0.2–2, using `FindClusters` function).

Cell cycle scoring (`CellCycleScoring` function) was performed using Seurat’s `cc.genes` and converted to mouse homologs using the MGI database.

This object containing all immune cells was used for major cell type (“coarse”) annotations. For cell type subset-specific annotations, the Seurat object was subsetted by monocytes, macrophages, neutrophils, and dendritic cells and the above Seurat and Harmony workflow was repeated prior to subtype (“fine”) annotations. Manual subtype annotation was performed using differential gene expression analysis (`FindAllMarkers` function, using MAST method and “experiment” as a `latent.vars`) and markers reported in the literature [11, 16–19, 23, 32].

### Metastatic Burden Quantification

Metastatic burden was summarized based on the data previously reported [15]. Total number of nodules was calculated by summing nodule counts across all categories (micro, intermediate, and macro) for each lung. Similarly, median nodule area was calculated by taking the median nodule area across all nodule categories for each lung.

### Principal Component Analysis of Cell Type Proportions

Principal component analysis was performed as previously described [13] on logit-transformed cell type proportions using `getTransformedProps` function from the `speckle` package. Transformed proportions were scaled according to cell type (not centered). PCA was performed using the `pca` function (svd method) from the `pcaMethods` package. PCA scores were extracted with the `score` function and loadings were extracted with the `loadings` function from the `pcaMethods` package.

### Cell Type Proportion Calculations

#### Tissue enrichment in fine cell types

The number of cells sequenced from each tissue type was uneven (Supp. Fig. S1A). Therefore, for each coarse cell type we equalized cell counts across tissues by random downsampling to match the tissue with the fewest cells (*n*/tissue = 1,186 for monocytes, 1,224 for macrophages, 1,011 for DCs, 40 for neutrophils). After downsampling, the proportion of each tissue in each fine cell type was calculated. We repeated this process of downsampling and calculating proportions 500 times, and averaged the proportions across all iterations and calculated standard deviations to quantify variability due to random downsampling.

#### Comparing cell type subset proportions across tissues

For the following sections, proportions of cell type subsets were calculated for each individual sample as a fraction of each subset’s corresponding major cell type (i.e. non-classical monocytes proportion was calculated as a fraction of all monocytes). Furthermore, samples with less than 15 total cells (within major cell type) were excluded from analyses. For visualization in figures, raw proportions were used. For statistical testing, we used logit-transformed cell type proportions using the `getTransformedProps` function in the `speckle` package. We fit a linear mixed-effects model for each cell type subset using the `limma` package, treating PDX model as a random effect to take into account variability between biological replicates. We used the `lmFit` function for fitting the models, `duplicateCorrelation` for estimating within-block (PDX model) correlation, and `eBayes` for empirical Bayes moderation to stabilize variance estimates. Benjamini-Hochberg correction was applied for multiple testing (within each major cell type).

#### Proportion in malignant vs. control tissues

To summarize the magnitude of differences between malignant tissues and their corresponding controls, cell type subset proportions per sample were averaged according to tissue type (primary, metastatic, control mammary gland or lung). We then calculated the ratio of the average proportion in primary tumor to mammary gland, or metastasis to control lung, and log-transformed the ratio for visualization. For statistical testing, we ran separate linear mixed-effects models for the primary-mammary gland and metastasis-control lung comparisons.

#### Proportion in primary tumors vs. metastases

To summarize the magnitude of differences between cell type subsets in primary tumor and metastatic lung samples, cell type subset proportions per sample were averaged according to tissue (excluding controls). Linear mixed effects models were fit as described above.

#### Changes in proportions between malignant tissues and control tissues

To summarize and compare the changes in cell type proportions between malignant tissues to those between control tissues, sample cell type subset proportions were again averaged according to tissue type. We then calculated the ratio of the average proportion in control mammary glands to control lung, and the ratio of the average proportion in primary tumor to metastases, and log-transformed each set of ratios for visualization. To distinguish between malignancy-specific and tissue-intrinsic compositional differences, we computed the following contrast: (Primary − Metastasis) − (Control MG – Control Lung).

#### Correlating fine cell type proportions and metastatic burden

Fine cell type proportions were calculated for each sample as a fraction of each major cell type. Spearman correlation was then run for all combinations of fine cell types and metastatic burden types (median nodule size, number of nodules, percent metastatic area) separately for primary tumor samples and lung samples. P-values were adjusted using the Benjamini-Hochberg method within groups according to tissue and cell type.

### Bray-Curtis Dissimilarity Calculations

To quantify inter-sample myeloid compositional heterogeneity, for each sample we first computed the logit-transformed proportion of all subsets within each major cell type (monocytes, macrophages, DCs, neutrophils) using the `getTransformedProps` function from the `speckle` package. Samples with fewer than 15 cells and control tissue samples were excluded from the analysis. Bray-Curtis dissimilarities were then computed on the resulting sample × cell proportion matrices using the `vegdist` function from the `vegan` package. To summarise our results, we then calculated the average dissimilarity for each sample to all other samples of the same tissue type, resulting in a mean dissimilarity metric for each sample ranging from 0–1, where 1 is most dissimilar and 0 is most similar. Differences in mean dissimilarity between primary tumor and metastatic lung samples were assessed using a Wilcoxon rank-sum test.

### Integrative Non-Negative Matrix Factorization (iNMF) and Network Module Analysis

iNMF was performed on the full Seurat object of all immune cells using the DECIPHER-seq-v2 package (adapted from [40]). Briefly, the original DECIPHER-seq code was modified to allow for compatibility with Rliger v2 and Seurat v5. DECIPHER-seq-v2 can be found at https://github.com/dasuperville/DECIPHER-seq-v2. The standard DECIPHER-seq workflow was followed as previously described [40]. “Sample” was defined as PDX model and tissue type (`pdx-tissue` in the object metadata), “batch” was defined as experiment (PDX1, 2, 3, 7, 9, 12), and “type” was defined as coarse cell type (monocyte, macrophage, dendritic cell, or neutrophil). Each cell type was present in at least 10 samples with at least 50 cells per sample. iNMF was performed using the `iNMF_ksweep` function on *k* = 2 through *k* = 40, with 20 repetitions.

To define activity programs for each cell type, we followed the standard DECIPHER-seq workflow for constructing phylogenetic trees, partitioning trees, and determining *k*. Notably, choosing an appropriate distance via `suggest_dist_thresh` for partitioning the phylogenetic tree did not result in appropriate partitioning, so we manually selected a threshold (by excluding the left tail of the pairwise patristic distance histogram plots) of 0.3, which fit all cell types. *K* values were as follows: *k* = 27 for monocytes, *k* = 26 for macrophages, *k* = 23 for DCs, *k* = 11 for neutrophils. Outlier programs representing rare cells were filtered out by DECIPHER-seq, resulting in 25 programs for monocytes, 25 programs for macrophages, 23 programs for DCs, and 8 programs for neutrophils.

Downstream analyses such as calculating expression score correlations across samples, determining marker genes for programs, constructing the DECIPHER-seq network, and performing GSEA on network modules were performed as described [40].

### Program-Metadata Associations

To identify programs that were significantly different between, or correlated with, particular metadata features, we created a statistical testing pipeline for testing every combination of program and metadata feature in each cell type. First, we added the H matrix (cells × NMF weights) from our NMF results to the metadata of each cell type-specific Seurat object. The metadata was grouped by sample number (unique to each animal, tissue, and sequencing batch), and the number of cells per sample was counted to exclude samples with fewer than 25 cells from the statistical analyses. All metadata-program tests were conducted separately on metastatic/control lung samples and primary tumor/control mammary gland samples (except for the tissue vs program tests, which inherently compared tissue types). Any samples that did not have a known value for a particular metadata feature were excluded from that particular program-metadata test. Statistical workflows differed slightly depending on the type of metadata feature, of which there were three: categorical metadata (e.g. tissue type), continuous metadata describing samples (e.g. metastatic burden), or continuous metadata describing cells (e.g. gene signature scores).

For categorical metadata (e.g. tissue associations, Fig. 3), we compared the mean program score across different metadata categories. To do this, we first collected summary statistics on each program-metadata comparison, including number of samples, mean and median program score, and standard deviation per metadata category, which were saved for downstream analysis. Any categories with fewer than 3 samples were excluded from the analysis because we would not be able to perform statistics on them. If, after removing metadata categories with fewer than 3 samples, we were left with less than two groups to compare, that program-metadata comparison was skipped. Next, we tested for homogeneity of variances (Levene’s test) and normality (Shapiro-Wilkes test). Our criteria for selecting the appropriate statistical test to compare means was as follows: metadata features with only two categories were either tested with a T-test (normally distributed, homogenous variances), T-test for unequal variances (normally distributed, non-homogenous variances), or Wilcoxon test (not normally distributed). Metadata features with more than two groups were either tested with one-way ANOVA test (normally distributed, homogenous variances), Welch’s ANOVA test (normally distributed, non-homogenous variances), or Kruskal-Wallis test (not normally distributed). For omnibus tests, we then ran post-hoc tests to determine which groups were statistically different and their adjusted p-values (Tukey test for one-way ANOVA, Games-Howell test for Welch’s ANOVA, Dunn test for Kruskal-Wallis).

For sample-level continuous metadata features (e.g. metastatic burden measurements, Fig. 3), we calculated the correlation between the sample metadata value and the average NMF program score across all cells per sample. As before, we calculated summary statistics (total number of samples, mean and median program score per sample, standard deviations). We next ran the Shapiro-Wilkes test on the distributions (mean program scores and metadata values) to determine if they were normally distributed. For normal distributions we performed Pearson correlation, for non-normal distributions we performed Spearman correlation. Notably, for correlations between metastatic burden and program score, control tissues were excluded as they came from tumor-free animals.

For cell-level continuous metadata (e.g. signature scores, Fig. 3 and 4), we aimed to calculate the correlation between cell metadata values and cell program weights. Here, we simply used the Shapiro test to determine normality and then performed either Spearman or Pearson correlations (as before). In cases where there were greater than 5,000 cells the Shapiro test could not be used, and we performed a Pearson correlation.

### Defining Batch-Associated Programs

Since PDX models are not consistently represented across batches, after performing NMF we wanted to make sure that none of the NMF programs were affected by batch. Batch effects were primarily seen between, rather than within, 10X chemistries. Therefore, we identified a few cross-chemistry batches that had substantial overlap in the PDX models that were sequenced (PDX3 and PDX7, and PDX1 and PDX12). We extracted all NMF scores for cells within PDXs that were present across these two sets of batches, and filtered out samples (and their cross-batch pairs) that had fewer than 25 cells. We calculated the average NMF score for each sample in each batch and repeated the statistical workflow described above (categorical workflow). Programs that were significantly different between both sets of batches were removed from subsequent analyses. Of note, we also performed this analysis between two batches sequenced with the same chemistry (PDX7 and PDX12), and found no statistically significant differences between batches in any programs.

### Defining Tissue-Associated Programs

Based on the results of the tissue vs program statistical tests (categorical workflow), we first identified programs that were significantly different between metastatic lung and primary tumor (padj *<* 0.05). Next, we identified whether any of those programs were significantly different between control lungs and control mammary glands. Based on the saved summary statistics (categorical workflow), we compared the magnitude of the means of each tissue. Programs were defined as: “metastatic lung” programs if they were significantly higher in metastatic lungs compared to primary tumors but not significantly different between control tissues; “primary tumor” programs if they were significantly higher in primary tumors compared to metastatic lungs but not significantly different between control tissues; “general lung” programs if they were significantly higher in metastatic lungs compared to primary tumors and significantly higher in control lungs compared to control mammary glands; “general mammary gland” if they were significantly higher in primary tumors compared to metastatic lungs and significantly higher in control mammary glands compared to control lungs.

### Defining Metastasis-Associated Programs

Metastasis-associated programs were defined if they were significantly correlated (*p <* 0.05) with percent metastatic burden, number of metastatic nodules, and/or median nodule size from the statistical workflow described above (continuous, sample level). Of note, programs that were significantly correlated with multiple metrics did so in a consistent direction (all positive or negative).

### Signature Scoring

All signature scores were calculated using the UCell package `AddModuleScore_UCell` function [66]. For the time-course analysis of metastasis-associated programs, the top 50 marker genes from each program (as determined by DECIPHER-seq) were selected for signature scoring.

### Time-course Analysis

Processed single-cell RNA-seq datasets from mouse E0771 [13], PyMT [11], 4T1 [11] and human HCI010 [12] models were obtained from the NCBI Gene Expression Omnibus database. When provided, processed cell type specific objects were used directly. Otherwise, count matrices were fully processed as described above and subset by major cell type (monocyte, macrophage, dendritic cell, neutrophil). The following workflow was performed for each cell type subset object in each dataset: First, cells were scored for metastasis-associated program signatures as described above. Then, the average signature score for each program was calculated for each sample (animal). Timepoint labels were standardized to ensure consistent naming across datasets. Notably in the E0771 dataset, samples from distal normal tissue, primary tumor tissue, and relapse-free tissue were excluded, as there were not comparable samples in the other datasets. Furthermore, “pre-metastatic” samples were designated as “early”, whereas “metastatic”, “metastatic core” and “metastatic invasive margin” were designated as late. The E0771 dataset lacked samples that were equivalent to “mid” timepoints from the other datasets. To determine whether program expression changed significantly over time, we performed a t-test comparing the mean program score for “early” vs “late” samples for each program, followed by Benjamini-Hochberg correction. Effect sizes and confidence intervals were calculated using the `effectsize` package `hedges_g` function.

### Calculating Percentile Rank of Program Genes

We compared the rank of each gene by weight in each program. NMF weighs each gene in the expression matrix according to its importance to each program, however, this weight value is not comparable across programs. Therefore, we standardized the gene weights by calculating the percentile rank of each gene for each program and then performed unsupervised hierarchical clustering on the ranked genes. Specifically, program genes were ordered by weight, and the `ecdf` function from the `stats` package was used to calculate the empirical cumulative distribution function (ECDF). The ECDF was then applied to the gene weights to calculate the percentile rank (between 0 and 1) for each gene based on weight.

### MDSC Marker Identification

To identify marker genes that distinguished MDSC Programs 8 vs 24, we identified the program which contributed most to each cell’s activity by ranking each program within each cell according to weight. Based on these rankings, we selected cells that had monocyte program 8 or monocyte program 24 as their top ranking program and defined them as the “top program 8/24 cells”. We then performed differential gene expression analysis using Seurat’s `FindMarkers` function between the top program 8 and top program 24 cells. Relevant DGE parameters include a log foldchange threshold of 0, only on genes expressed in at least 0.1% of cells, the ‘MAST’ test, and ‘experiment’ as a latent variable. Gene set enrichment analysis was done on the resulting DGE list sorted by average log_2_FC with the Reactome gene sets using the `fgsea` package’s `fgsea` function (min.size = 4).

### SCENIC and Regulon-NMF Correlation Analysis

pySCENIC v0.12.1 was run using the apptainer image on count matrices from each major cell type object separately, and for the 4T1 monocyte time-course dataset [11]. We used the mm10 refseq_r80 rankings database, the allTFs_mm transcription factor list, and the v10 motif2TF annotation (mgi). To calculate correlation between regulon activity and NMF program activity, we used the AUC matrix (regulon activity per cell) and NMF H matrix (NMF weight per cell). For each program-regulon pair, we tested for normality using the Shapiro test when possible (*<* 5000 cells). For normal distributions we calculated Spearman correlation. For non-normal distributions or cases of *>* 5000 cells, we calculated Pearson correlation. P-values were adjusted for multiple comparisons (for each major cell type) using the Benjamini-Hochberg method.

## Acknowledgements

We thank C.M.M. Diadhiou, A. Abbasi, S. Hasnain, E. Atamaniuc, L. Awni, J.V. Lee, A. Nanjaraj, V. Sitarama, V. Guchshina, E. Weigert, A.L. De Souza, Ç. Yaha, N. Singh, S.P.C. Wagner for technical assistance. We thank S. Tikoo for fruitful discussions and input.

Sequencing was performed at the UCSF CAT, supported by UCSF PBBR, RRP IMIA, and NIH 1S10OD028511-01 grants.

We acknowledge the PFCC (RRID:*SCR_018206*) supported in part by Grant NIH P30 DK063720 and by the NIH S10 Instrumentation Grant S10 1S10OD021822-01.

This study was supported by funds from the European Molecular Biology Organization (EMBO) Long-term Post-doctoral Fellowship (EMBO ALTF 159-2017, to JW); a Program for Breakthrough Biomedical Research Award (to JW); an ImmunoX Bakar Trainee Momentum Award (to JW), a Helen Diller Family Comprehensive Cancer Center Breast Oncology Program Award (to JW), Vienna Science and Technology Foundation (WWTF-LS23-067, to JW), an NIH F31 award (F31CA284749, to DAS), R01 CA056721 (to ZW), CDMRP DoD W81XWH-21-1-0774 (to AG), The Mark Foundation (to AG), Bechtle Family Foundation (to AG), and Subramanian Breast Cancer Support Fund (to AG).

## Author contributions

JW conceptualized and supervised the study. JW performed experiments, and generated scRNA-Seq data with the help of DAS and EC. DAS analyzed scRNA-seq data with the help of CM, YZ and AR. AR, CM, IB, and AJC provided intellectual input and critical feedback. Funding was acquired by JW, AG, DAS, and ZW. DAS and JW wrote the manuscript with the input from all authors.

## Supplementary Figures

**Figure S1.**
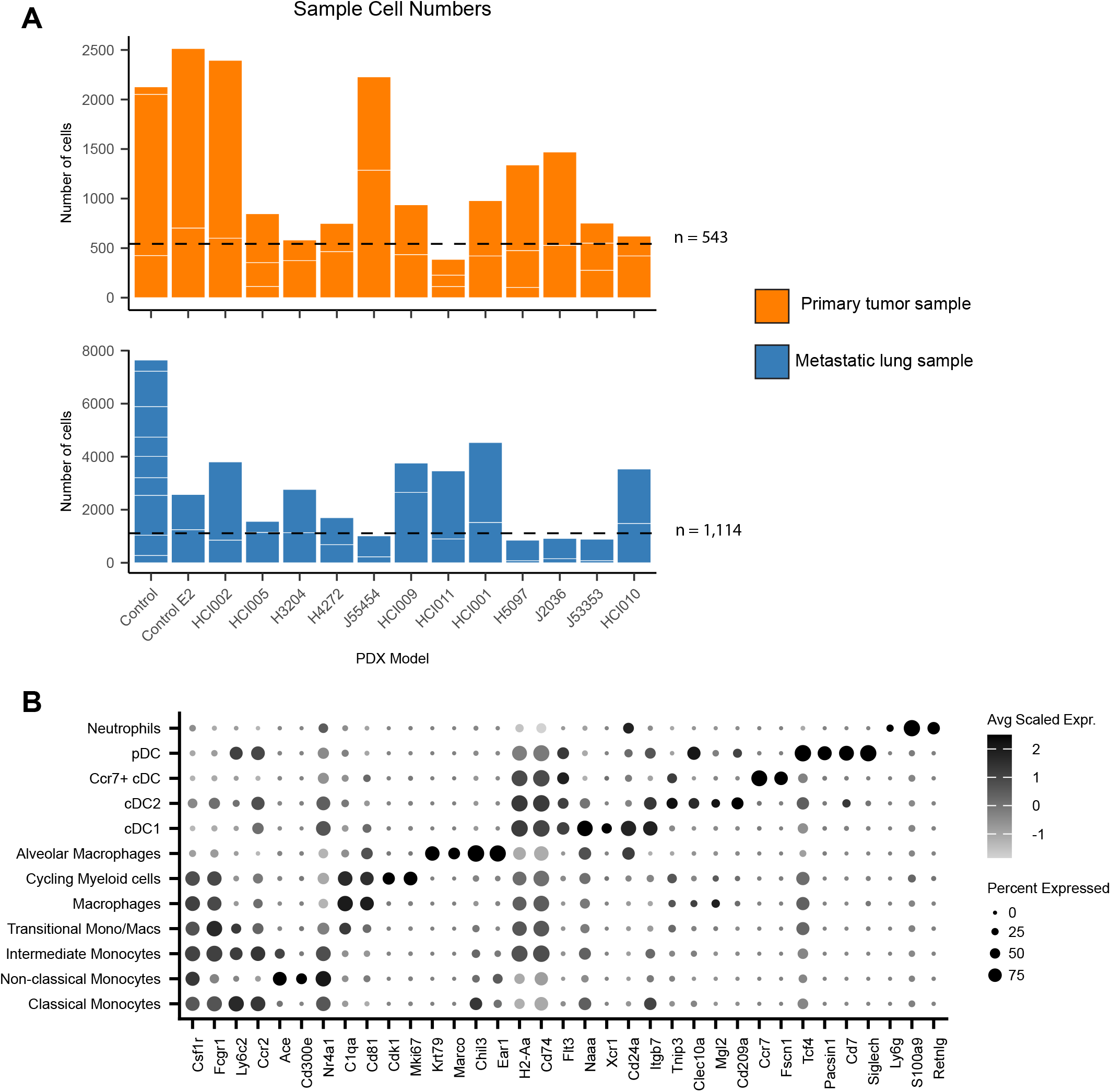
**(A)** Total number of singlets passing QC for each sample. Samples are stacked according to PDX and tissue. Dotted line indicates average number of QC+ cells per sample for each tissue (543 cells for tumor/mammary gland samples, 1114 for control/metastatic lung samples). **(B)** Scaled average expression of markers used for annotation of major immune cell types.

**Figure S2.**
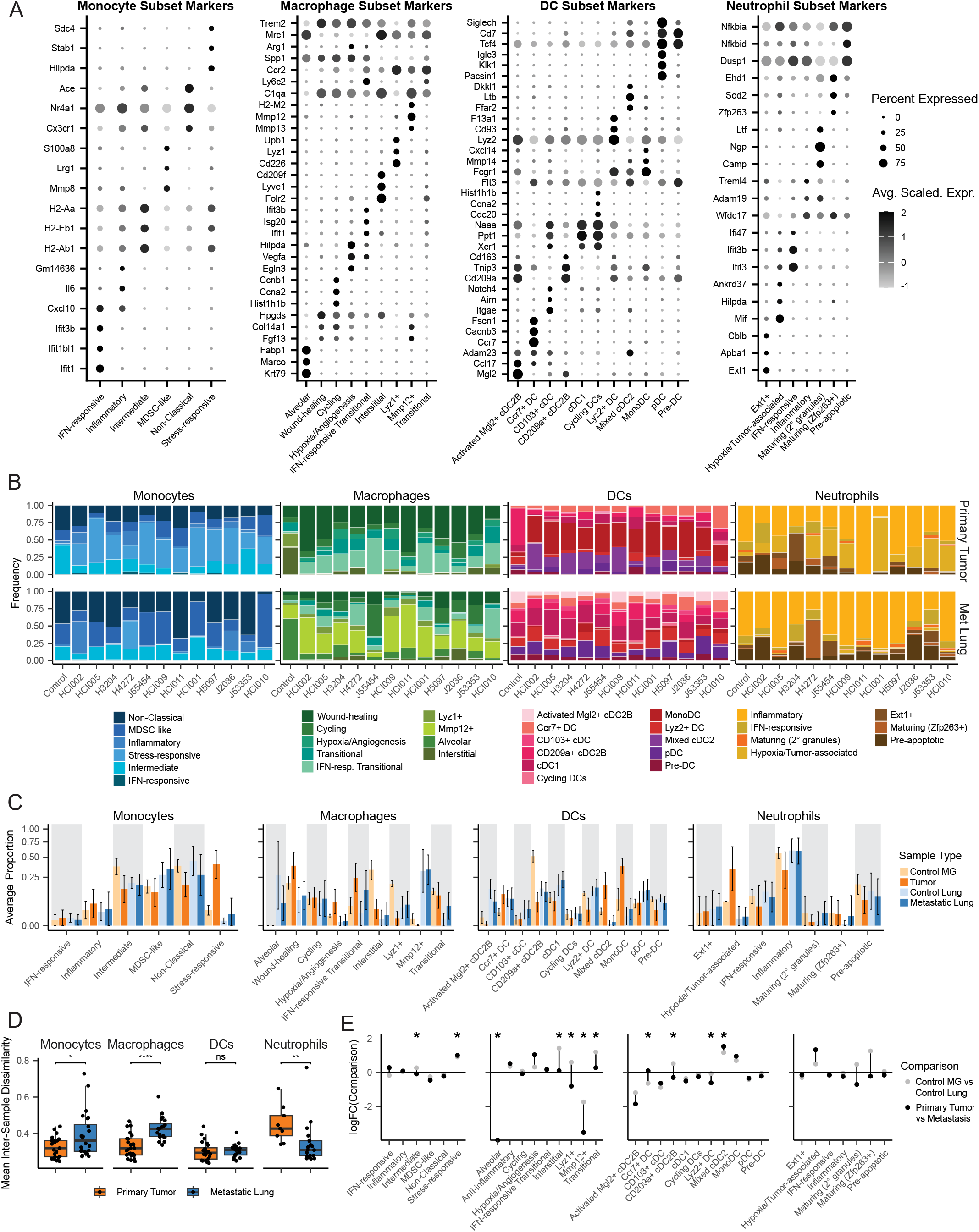
**(A)** Average scaled expression of select top markers of each cell type subset identified by subclustering of monocytes, macrophages, dendritic cells and neutrophils. **(B)** Proportion of cell type subsets per PDX model and tissue. Cell type subset proportion calculated as a frequency of major cell type. **(C)** Average proportions of cell type subsets across samples. Subset proportions were calculated as a fraction of each major cell type within individual samples, averaging samples according to tissue type. Error bars indicate standard deviation. **(D)** Mean inter-sample Bray-Curtis dissimilarity (within tissue type) for each major myeloid cell subset. Each point represents one sample’s mean Bray-Curtis dissimilarity to all other samples of the same tissue type. Bray-Curtis dissimilarity was calculated based on logit-transformed subset proportions for each major myeloid cell type. Statistical significance of mean dissimilarity among primary tumors and metastatic lungs was calculated using a Wilcoxon rank-sum test. NS p > 0.5; * p < 0.05; ** p < 0.01, **** p < 0.0001. **(E)** Log fold-change of average proportions between control tissue samples (Control MG vs Control Lung) and malignant tissue samples (Primary Tumor vs Metastasis). * p < 0.05, nested linear mixed effects model with PDX as a random effect, comparing the change between control tissues to the change between malignant tissue samples, followed by empirical Bayes moderation and Benjamini-Hochberg multiple testing correction.

**Figure S3.**
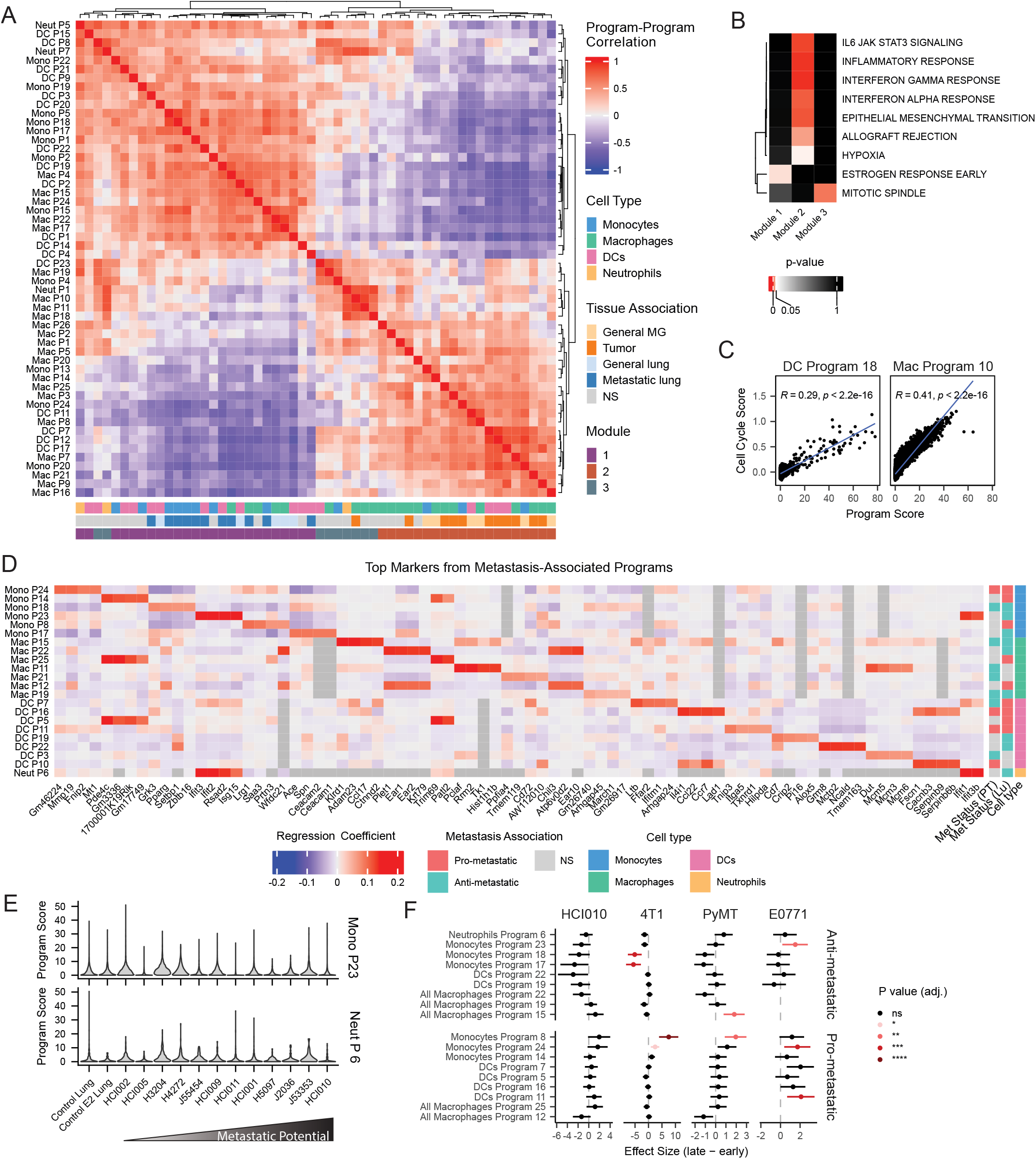
**(A)** Pairwise correlation between all NMF programs that passed filtering and were classified into a module by DECIPHER-seq. Annotations indicate program cell type, tissue status, and module. **(B)** Correlation between cell cycle score calculated in Seurat and program score for DC Program 18 and Macrophage Program 10 “cell cycle associated” programs. (C) Gene set enrichment analysis of DECIPHER-seq modules using Hallmark gene sets. All significant gene sets for each module are shown. Gene sets were defined as significant if they were significantly enriched in three or more NMF programs per module. **(D)** Top marker genes for each metastasis-associated NMF program. Marker genes and regression coefficients were calculated as described in Murrow et al. **(E)** Distribution of IFN program scores across PDX lungs. Tumor models are ordered according to metastatic potential (left to right, low to high). **(F)** Effect sizes and 95% CI of metastasic program signature scores between early and late timepoints. Programs are colored according to significance (Benjamini-hochberg corrected).

**Figure S4.**
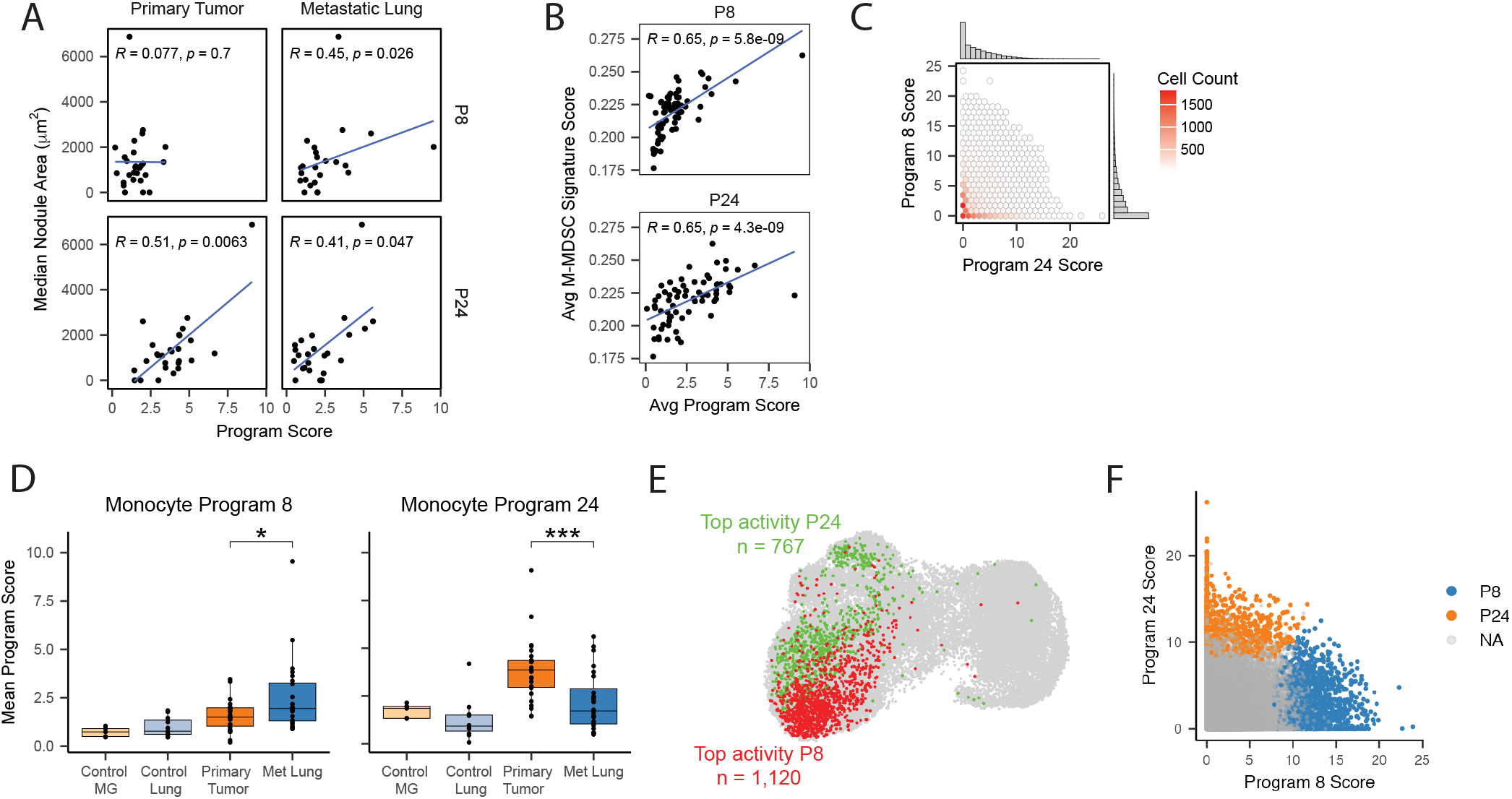
**(A)** Spearman correlation between average program 8 and 24 score across all monocytes per sample and sample median nodule size, split by sample tissue type. **(B)** Spearman correlation between average M-MDSC signature score (*Alshetaiwi et al*.) and average NMF program score per sample across all monocytes. **(C)** Distribution of program 8 and 24 scores across monocytes (cells filtered for NMF weight > 0 for either program). Color indicates cell frequency, marginal histograms highlight individual program distributions. **(D)** Comparison of average program score per sample across all tissues for monocyte programs 8 and 24. Statistical testing only for primary tumor vs metastatic lung. T-test for Program 8, Wilcox test for Program 24, * = p < 0.05, *** = p < 0.001. **(E)** UMAP highlighting cells selected for differential gene expression analysis based on program 8 (P8, green) or program 24 (P24, red) activity. Program activity was determined by ranking NMF programs by weight within cells, and identifying the top ranking program with each cell. **(F)** Scatter plot of program 8 and program 24 scores across all monocytes. Top program 8 (P8) and program 24 (P24) cells are highlighted in blue and orange, respectively. Cells not included in differential gene expression analysis are in grey (“NA”).

**Figure S5.**
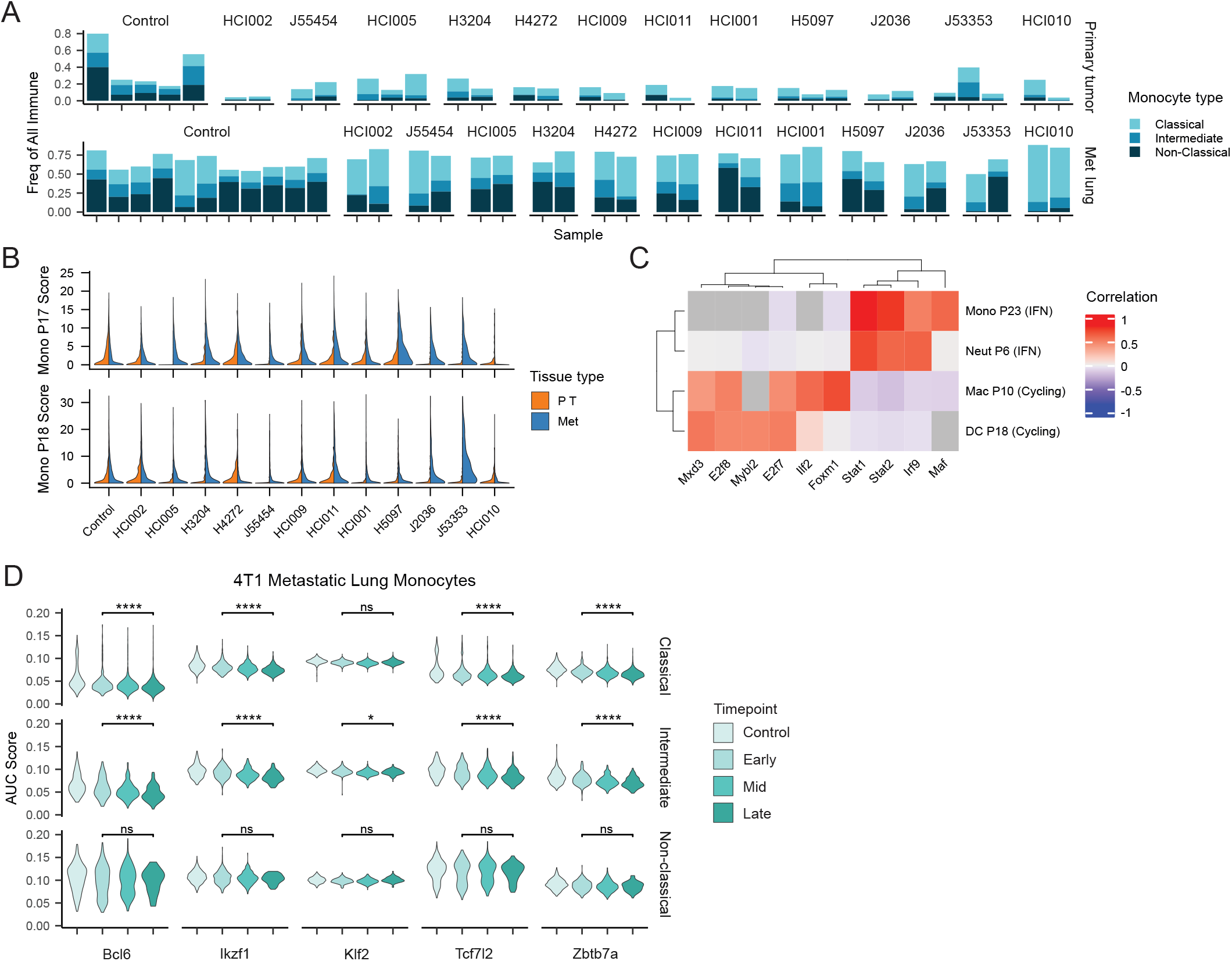
**(A)** Frequency of classical, intermediate, and non-classical monocytes of all immune cells per sample. Bars represent individual samples and are grouped according to PDX model and tissue. **(B)** Distribution of program 17 and program 18 scores across all monocytes per PDX model and tissue. **(C)** Heatmap of spearman correlations for the top correlated regulons for monocyte program 23, neutrophil program 6, macrophage program 10, and DC program 18. Proposed function in parentheses next to each program. **(D)** Distribution of activity scores for regulons involved in classical to non-classical monocyte differentiation across all monocytes in 4T1 lungs. Distribution of regulon acitivty are shown for early, mid, and late timepoints as well as control samples. Wilcoxon rank-sum test (early > late), adjusted for multiple comparisons using the Benjamini-Hochberg method. * p < 0.05, **** p < 1e-4, ns not significant.

## Notes

### Competing Interest Statement

The authors have declared no competing interest.

